# SAME: Topology-flexible transforms enable robust integration of multimodal spatial omics

**DOI:** 10.1101/2025.07.12.664419

**Authors:** Aditya Pratapa, Siavash Mansouri, Nadezhda Nikulina, Bruno Matuck, Marc A. Schneider, Kevin Matthew Byrd, Rajkumar Savai, Purushothama Rao Tata, Rohit Singh

## Abstract

Spatial omics technologies provide complementary and layered molecular insights that span proteins, transcripts, and metabolites. However, aligning and integrating these modalities across serial tissue sections remains a computational challenge. Existing alignment methods are primarily unimodal and assume preserved topology, often failing with tissue distortions like tears, folds, or anatomical changes. Here, we present SAME (Spatial Alignment of Multimodal Expression) that introduces space-tearing transforms, a framework for controlling localized topological disruptions during cross-sectional alignment. Using integer linear programming to maximize cell type matches across the modalities, we enhance cell-type alignment accuracy by 20% compared to existing methods while preserving biologically meaningful spatial relationships. Applied to protein-RNA integration in healthy tongue tissue and lung adenocarcinoma, SAME revealed cryptic immune subpopulations that were otherwise missed by RNA-only or protein-only classification. In a separate lung adenocarcinoma study, we assayed and integrated protein and metabolomic profiles, uncovering localized mevalonic acid upregulation specifically within tumor-macrophage spatial niches and identifying targeted metabolic crosstalk invisible to single-modality approaches. SAME enables unprecedented experimental designs that leverage each modality independently while computationally recovering cross-modal spatial structure, unlocking multimodal discoveries in complex tissues.

## Introduction

Understanding spatial niches within complex and dynamic tissues like tumor immune microenvironments and regenerating tissues requires characterizing cells, subtypes, and cell states across diverse molecular perspectives. Emerging spatial omics technologies now provide this capability, but often require multiple modalities [1, 2]. For example, while current multiplexed immunofluorescence panels reliably identify cell types like B and T cells based on protein marker, spatial transcriptomics is often better suited to reveal gene expression programs related to signaling pathway activity or metabolic state, which are not always accessible at the protein level [3, 4, 5, 6, 7, 8]. Additionally, because some profiling methods are destructive, different modalities must often be applied to serial or near serial sections. Integrating data across modalities and serial sections thus demands the development of new computational approaches [7, 8].

Integrating diverse multimodal spatial data presents significant computational hurdles. The fundamental challenge is developing spatial transforms expressive enough to capture both biological variability and experimental artifacts while remaining computationally tractable. Broadly, spatial integration techniques enumerate potential correspondences of spots across slides, then compute the optimal spatial transformation that preserves the geometric relationships of these matched cells [9, 10, 11]. Unlike previous transcriptome-only settings, in our multimodal integration the correspondence has to be estimated across disparate measurement modalities, exacerbating the challenge. The earliest spatial integration efforts used simple rigid or affine (i.e., linear) transformations to align spots across slides [10]. More recent methods employ diffeomorphic (topology-preserving) transformations [12, 11, 13] that can handle limited tissue deformations where the topology is preserved. However, tissue topology itself can vary between serial sections, e.g., when a vessel branches in one section but not another. Topological changes such as tears and folds are also introduced by the physical sectioning and assaying process. These changes violate the assumptions of linear or diffeomorphic methods, creating a critical gap in spatial integration capabilities.

The need for topological flexibility is particularly acute in spatial biology applications. Spatial alignment methods have borrowed heavily from medical image registration, where minor topological violations could be treated as imaging artifacts to be ignored. However, contemporary spatial omics employ unbiased, genome-wide assays that capture rare but disease-critical cellular niches: small populations of drug-resistant cells, immune-privileged sanctuaries, or metabolically distinct tumor regions. Rigid adherence to topology-preserving transforms risks can misalign small spatial niches, resulting in inaccurate or incomplete characterization of cell-cell interactions. Conversely, arbitrary topological violations lead to nonsensical mappings where distant tissue regions appear adjacent, confounding biological interpretation. The field has lacked principled integration methods to permit controlled, localized topological flexibility.

We introduce SAME (Spatial Alignment of Multimodal Expression) to integrate diverse spatial modalities across serial or near-serial tissue sections with unprecedented spatial flexibility (Fig. 1). The key conceptual advance of SAME lies in introducing a powerful category of spatial transforms to the integration problem. *Space-tearing transforms*, as we term them^†^, allow controlled topological violations within a broadly diffeomorphic alignment through a constrained optimization formulation. This permits localized topological disruptions where biology demands while maintaining overall tissue coherence. To compute space-tearing transforms, we introduce an efficient geometric measure of local topological change through triangle inversion counts. We then formulate an integer linear program (ILP) to compute the optimal transformation that balances triangle inversions with overall correspondence.

**Figure 1.**
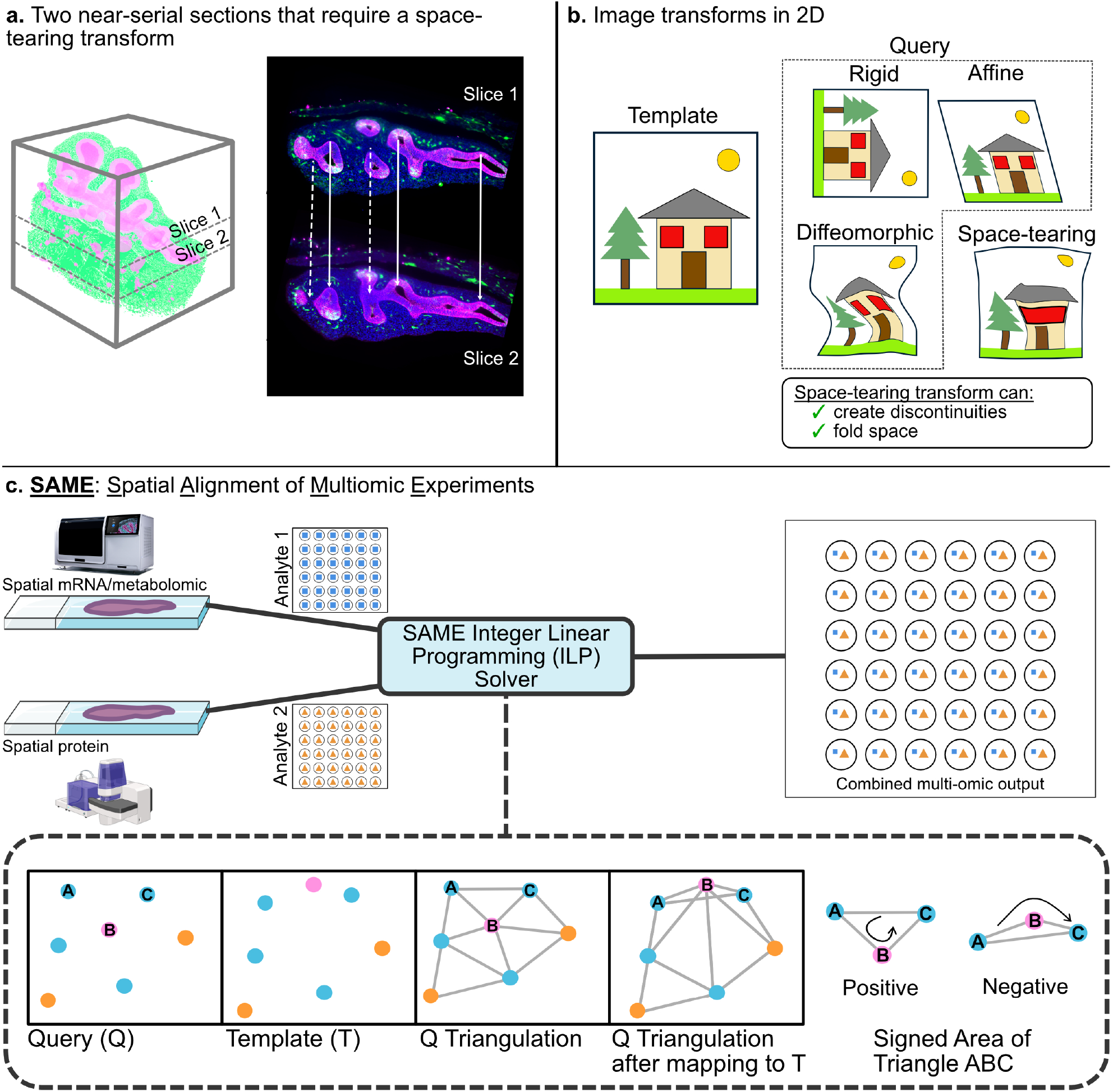
Overview of SAME workflow. a. Two near serial sections in a developing lung tissue that will require a space-tearing transform to match the two sections. Solid lines represent areas that can be aligned with standard affine or diffeomorphic transformations, while dotted lines denote regions needing a space-tearing transformation. b. Types of spatial transforms needed to transform the query image to template. Existing methods use rigid, affine, or diffeomorphic transforms that suffice if the topology is preserved. However, if the topology changes (here, two windows merged into one), a space-tearing transform is needed. c. SAME starts with a pair of two different spatial modalities and solves an integer-linear programming (ILP) problem to maximize cell-type matches across the modalities while simultaneously minimizing geometric distortions by using signed triangle area as the proxy for space-tears.

The other critical innovation in SAME is establishing cross-modal correspondences by probabilistically matching phenotypes (i.e., cell types) rather than comparing raw molecular counts. This sidesteps the complexities inherent in aligning analytes across modalities; for instance, transcript and protein measurements of a gene often correlate poorly. SAME computes phenotypes within each modality by leveraging protein markers, transcriptional programs, or tissue morphological features obtained through pre-trained foundation models for H&E images. We demonstrate SAME’s efficacy through multiple applications. First, on near-serial heart sections with complex tissue architecture and challenging, non-linear distortions, SAME outperforms existing approaches by over 20 percent. We then leverage SAME’s unique multimodal capabilities to integrate spatial protein data with spatial transcriptomics, revealing spatially distinct T cell functional states across healthy and tumor tissues that were undetectable in RNA-only profiling due to lymphocyte dropout. Finally, we pioneer the integration of spatial proteomics with metabolomics: in lung adenocarcinoma, we uncovered localized mevalonic acid upregulation within tumor-macrophage niches, a metabolic crosstalk invisible to single-modality approaches. SAME enables unprecedented experimental designs, providing the spatial biology community with a robust framework that maximizes information acquisition and integration across modalities.

## Results

### SAME: Spatial Alignment of Multiomic Experiments

SAME addresses multimodal alignment between two near-serial sections: a template slide *T* and a query slide *Q* to be aligned to *T*. Common use cases include aligning spatial protein data (e.g., from PCF) with spatial transcriptomics (e.g., Xenium, Visium, MERFISH) or spatial metabolomics (e.g., MALDI-MSI). We start by coarsely registering *Q* to *T* using standard rigid, affine, or diffeomorphic transformations based on nuclear (DAPI) or H&E staining channels [14, 15, 16]. This establishes a baseline correspondence while maintaining tissue continuity. In what follows, we use *T* ‘s coordinate frame as the reference frame. However, substantial spatial mismatches often remain even after this step due to biological variations across slides and physical artifacts like tears, distortions and folds (Fig. 1a–b, Fig. S1).

To support alignment across modalities, SAME first performs phenotype assignment (i.e., cell typing) on cells in each slide. The precision of this phenotyping depends on the modality: protein-based assays like PCF can yield accurate cell type calls using known markers; in transcriptomics, probabilistic methods such as RCTD [17] and cell2location [18] can infer likely identities; in metabolomics, we estimate phenotypes by extracting features from the corresponding H&E image, leveraging pathology-based pre-trained foundation models (Methods) [19, 20].

Crucially, SAME does not require perfect phenotyping. The algorithm uses these phenotypic estimates—whether binary labels or probabilistic distributions—to score candidate matches between cells across slides (Fig. 1c). Our goal is to match cells that are both phenotypically similar and spatially proximate. Spatial proximity here is defined using a *k*-nearest neighbor graph or a fixed-radius search in *T* ‘s coordinate frame established during coarse alignment. However, matching cells based only on local proximity and phenotype can result in globally incoherent mappings, i.e., alignments that preserve cell type identity but disrupt the overall tissue architecture. To minimize unnecessary topological disruption SAME introduces geometric regularization into the alignment formulation, selectively allowing local topological changes only where the data justify them. SAME constructs a Delaunay triangulation over the query slide’s cell centroids, capturing the tissue’s geometric mesh (Fig. 1c). It monitors triangle orientation by computing each triangle’s signed area. The sign of the area reflects whether the triangle’s vertices are ordered clockwise or counterclockwise, and a sign flip after mapping indicates a topological violation, serving as a geometric proxy for space tears. Limiting topological changes in *Q* to spatially-proximate cells in *T* ensures such changes remain local (typically less than 100 *µ*m). Increasing the search radius allows greater topological distortions, but also results in increased run time. To penalize triangle flips while maximizing valid cell-type matches, SAME formulates an integer linear program (ILP). An ILP formulates an optimization task over discrete variables subject to linear constraints, making it well-suited for assignment problems like ours where each query cell must map to exactly one template cell. Our ILP formulation includes binary variables for each potential cell-cell correspondence, with constraints ensuring one-to-one matching and an objective function that rewards phenotype matches while penalizing triangle orientation flips. This tunable framework allows localized space-tearing where needed while preserving overall geometric integrity.

### SAME maintains tissue geometry under both smooth and discontinuous distortion

Before applying SAME on experimental data, we assessed it using synthetic spatial data where we could control distortion types and evaluate the need for space-tearing transforms against simpler alternatives. We created two synthetic slices with a template slice *T* containing 144 cells in a 12 *×* 12 grid, with cell types assigned in an alternating pattern between two classes (purple and orange). To generate query slices *Q*, we applied two types of distortion to the template: a diffeomorphic warp using a radial basis function kernel, and a space-tearing warp created by superimposing random noise onto the diffeomorphic warp in the bottom right corner (Fig. 2b-c).

**Figure 2.**
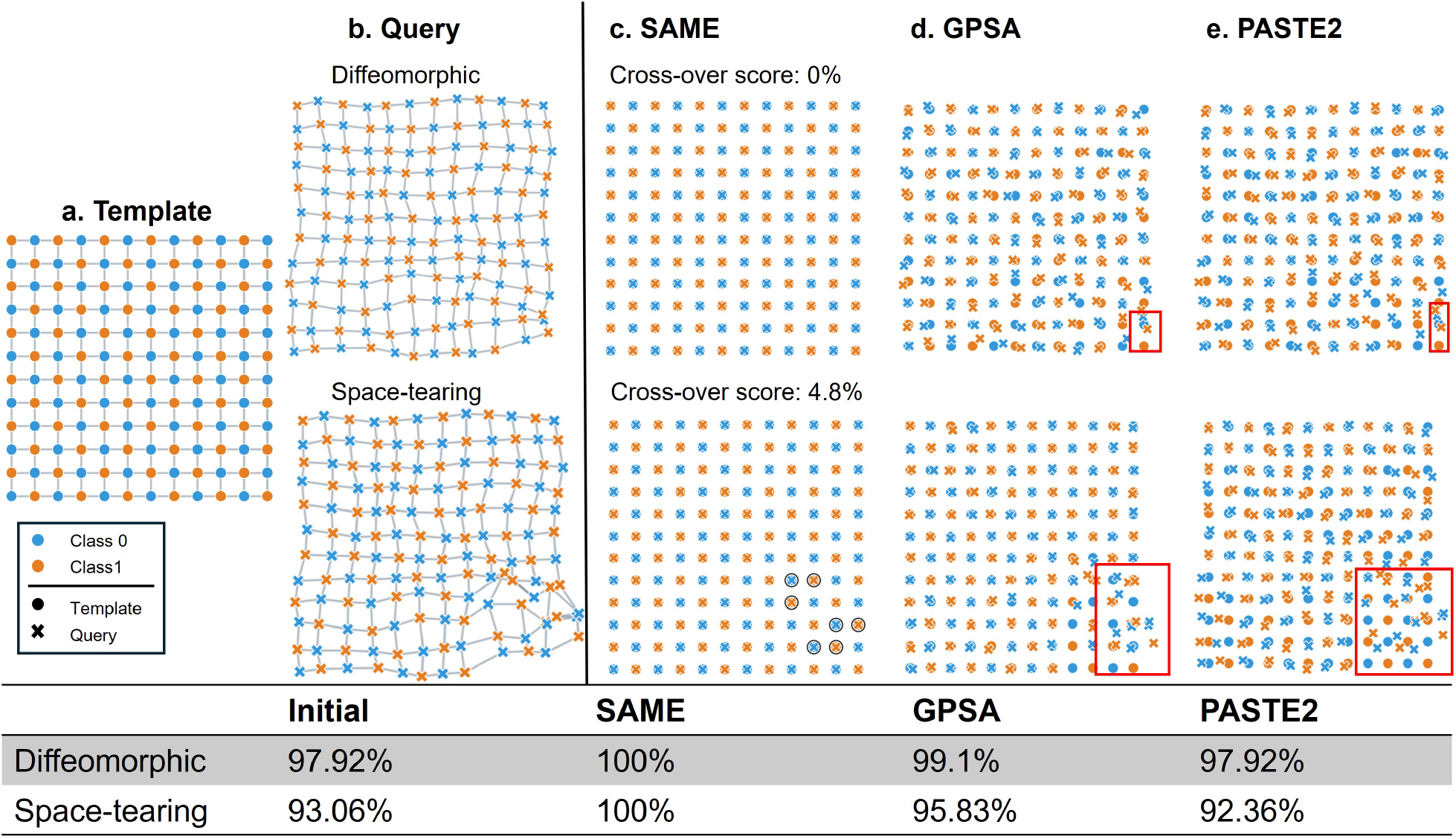
Performance of SAME under synthetic warps. a. Template grid showing the regular 12 × 12 grid configuration for 144 cells (marked by ‘o’)corresponding to two classes (blue and orange). b. Query slice cells (marked by ‘x’) obtained by applying a synthetic warp that is diffeomorphic (top row) and a space-tearing (bottom row) transform. c. Performance of SAME, d. GPSA, and e. PASTE2 under the synthetic warps.

We compared SAME against two state-of-the-art spatial transcriptomic alignment methods, GPSA and PASTE2, in aligning each distorted query slice *Q* back to the original template *T*. Exceeding other methods, SAME achieved perfect alignment (100% cell type matches) in both scenarios with minimal topology violations. For the diffeomorphic warp, SAME preserved local topology completely, with 0% triangle orientation violations. For the space-tearing warp, SAME maintained robust performance with only 4.8% triangle violations while still achieving perfect correspondence. Thus, SAME’s optimization model could prioritize only a limited, key set of local distortions, demonstrating SAME’s ability to allow controlled topological changes.

PASTE2, whose optimal transport formulation is designed for affine alignments, struggled with these non-linear distortions. GPSA, designed for diffeomorphic transformations, performed better than PASTE2 and achieved 99.1% accuracy in the case of diffeomorphic warp. For the space-tearing warp, GPSA improved from the initial 93% to 95.8% accuracy but remained limited due to its topology-preserving constraints. These results demonstrate that SAME effectively handles both smooth diffeomorphic distortions and irregular space-tearing distortions. This versatility makes SAME suitable for real biological samples, which frequently exhibit complex combinations of such distortion types.

### SAME outperforms existing methods in spatial transcriptomic integration

We next analyzed an *in-situ* sequencing (ISS) dataset from a 6.5 post-conception week human embryonic heart [21]. This dataset provides a valuable test case for alignment methods as it represents a real biological tissue with defined anatomical structures and cell type heterogeneity. This unimodal alignment scenario should favor existing expression-based alignment methods like PASTE2 and GPSA, since both slides measure transcriptomic readouts. We used the eight cell types defined from ISS expression values in the original publication to evaluate alignment accuracy (Fig.3a–b). Initial DAPI-based image alignment using Valis [16] yielded 57% cell-type matching accuracy (‘Initial’ in Fig. 3c; Fig. S1h) between query *Q* and template *T* slices, establishing our baseline.

**Figure 3.**
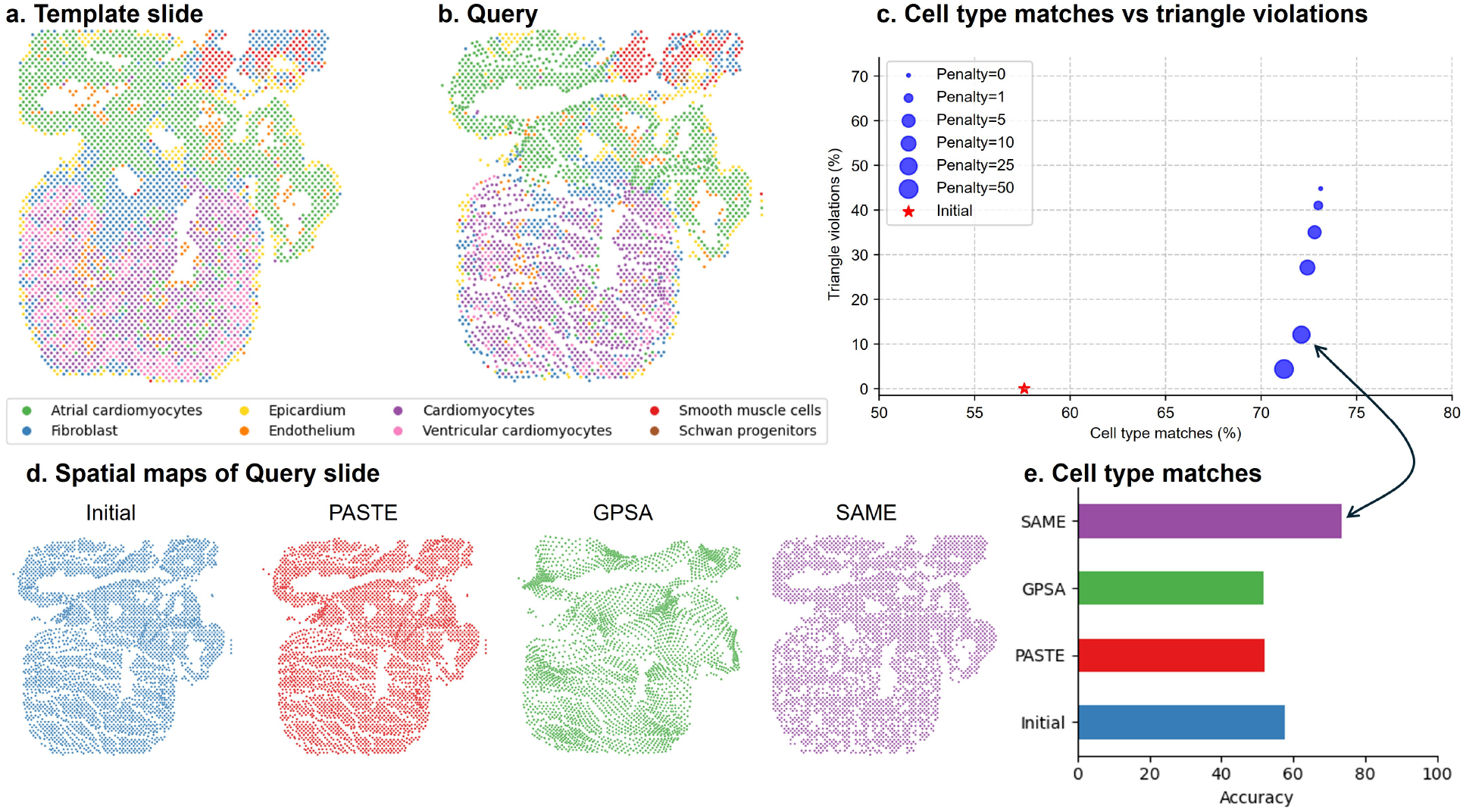
Serial section alignment of spatial transcriptomics (human embryonic heart). a. Cell-types identified in the Template and b. Query slices using smFISH-based profiling. c. Analysis of variation in the alignment accuracy compared to the cross-over violations. d. Spatial maps prior to alignment (Initial; blue) and after aligning with PASTE (red), GPSA (green), and SAME (purple). e. Cell-type accuracy results for the three methods considered.

SAME demonstrated tunable control over the accuracy–topology trade-off. When SAME was configured to allow for maximum topological flexibility by setting the penalty for triangle area sign inversions to zero, it achieved its highest cell-type matching accuracy of 73% for correspondences identified within the specified search radius. This peak accuracy, however, was accompanied by a substantial number of local distortions, corresponding to approximately 80% triangle orientation violations (Fig. 3c., penalty=0 point). To balance accurate cell-type mapping with the preservation of tissue topology, we systematically explored the effect of increasing this penalty term (Fig. 3c, Fig. S4). As the penalty for such orientation violations was increased, SAME progressively reduced the percentage of triangle violations. For instance, a high penalty (e.g., penalty=50) minimized these violations to less than 5%, while maintaining a high cell-type matching accuracy of 71.2%. This demonstrates SAME’s tunable capability, allowing users to prioritize either achieving maximal cell-type linkage in the presence of significant dissimilarities (by permitting more space warps) or enforcing stricter geometric consistency, potentially at a minor cost to the raw matching score. The observation that even with maximal allowance for distortions (penalty=0), the cell-type matching accuracy for this dataset plateaued at 73%, underscores that inherent biological differences between these two near-serial sections likely impose a fundamental limit on achieving perfect one-to-one correspondence.

SAME significantly outperformed existing methods even in this single-modality setting that is favorable to them (Fig. 3e). SAME achieved a final cell type matching accuracy of approximately 72% (penalty=25) with 12% of space-tearing matches. This significantly outperformed the initial DAPI-based alignment (57%), as well as other methods designed for similar data, such as PASTE2 (52% accuracy) and GPSA (51% accuracy). The qualitative results (Fig. 3d) further show that SAME produced a more coherent and anatomically plausible alignment of the query slice to the template structure compared to the other methods. These findings highlight SAME’s robust performance in aligning challenging real-world biological sections by optimizing for cell-type correspondence while actively minimizing topological inconsistencies, even in single-modality datasets where expression similarity is the primary guide.

### SAME demonstrates T cell functional programs are shaped by spatial niches

Mucosal linings are known to harbor specialized immune microenvironments in which T cells adopt spatially distinct functional states [22, 23, 24, 25]. However, such tissue-resident lymphocyte populations remain incompletely characterized, largely due to the limitations in current spatial profiling technologies. RNA-based platforms often fail to detect lymphoid cells reliably, as key T cell markers such as CD3E, CD4, and CD8A are expressed at low levels and subject to high dropouts [26, 27]. This issue is further compounded in resident T cells, which frequently downregulate conventional lymphoid markers while acquiring context-specific transcriptional programs.

Protein-based spatial profiling offers more reliable immune cell-type identification through surface markers but lacks transcriptional depth. In contrast, transcriptomic platforms offer a rich readout of gene expression states but cannot reliably resolve sparse immune populations, especially in complex tissues. These modality-specific limitations pose a challenge for understanding how microanatomical location shapes immune function.

To address this, we applied SAME to two near-serial sections (10–15 *µ*m apart) of healthy dorsal tongue tissue, selected for its well-defined epithelial organization and resident immune populations [25]. One section was profiled using the Akoya PhenoCycler-Fusion (PCF) system with a panel of 44 antibodies targeting lineage and functional markers (Fig. 4a, Fig. S5). The adjacent section was profiled using the Vizgen MERSCOPE platform with a panel of 300+ mRNA targets. This dual-modality design allowed for high-confidence cell-type identification through protein markers while capturing broader transcriptional states in spatially matched cells.

**Figure 4.**
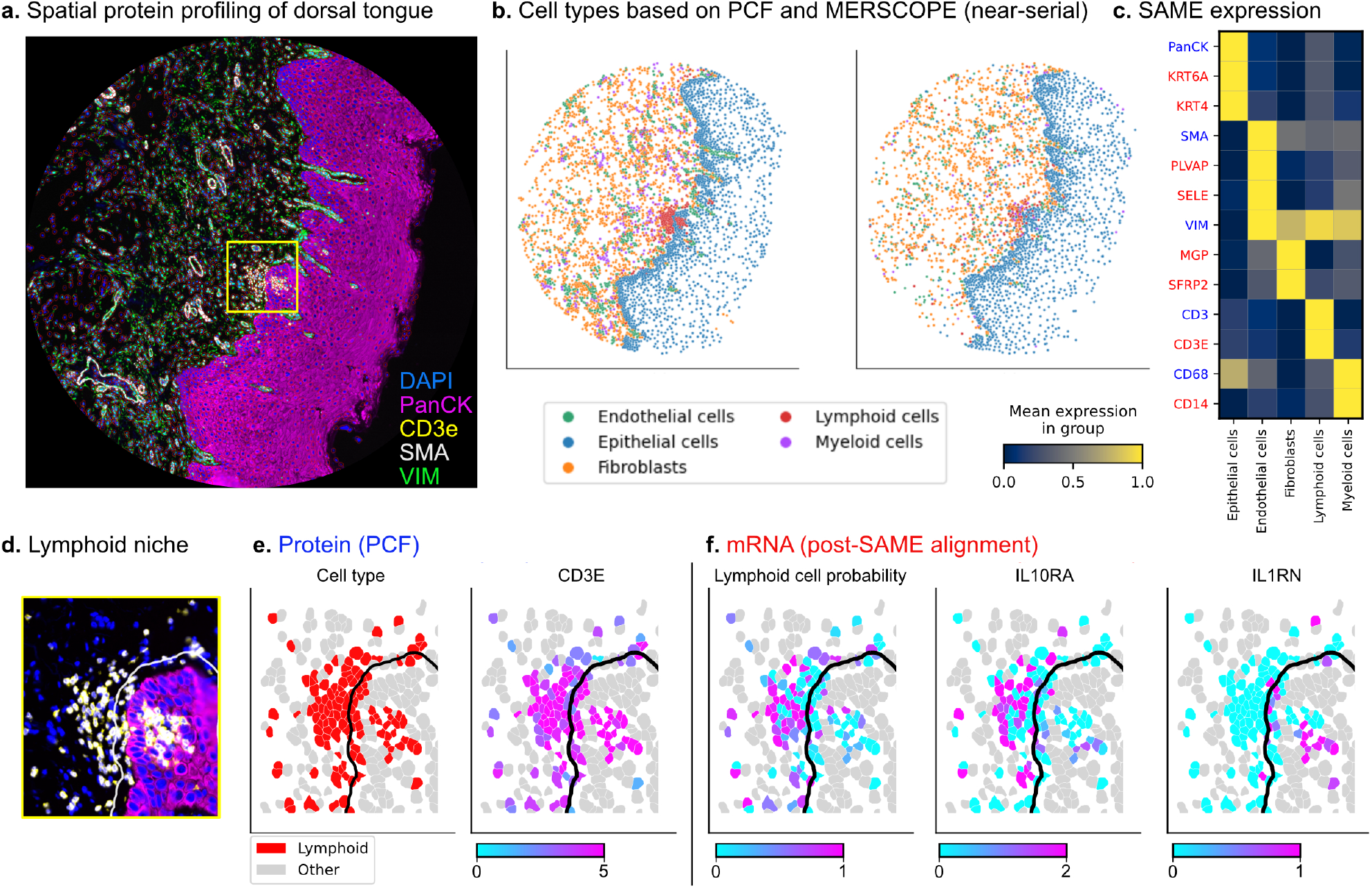
SAME integration of protein and mRNA profiling on near-serial sections reveals a lymphoid niche. a. Immunofluorescence profiling of the dorsal tongue using PCF showing key cell-type defining antibodies. b. Cell types are defined based on PCF (left) and MERFISH features (right) on near-serial sections. c. Heatmap depicting SAME-derived combined expression with protein markers in blue and transferred mRNA (red). d. Lymphoid cell niche (CD3E, yellow) showing subpopulations clustered within and around a structured epithelial region marked by PanCK (magenta). The boundary (white) outlines the epithelial layer and immediately surrounding cells (within 15 µm), defining the juxta-epithelial region. e. Protein-based annotation identifying CD3E-enriched lymphoid cells within and outside the boundary (black). f. SAME-aligned mRNA data show spatially distinct expression patterns, with intraepithelial lymphoid cells upregulating IL1RN, while IL10RA is enriched in laminar cells.

Post-segmentation and quality control yielded 4,671 cells from the protein dataset and 3,608 from the mRNA dataset. The reduced cell count on the RNA side reflects the limited transcript capture per cell, particularly for lowly expressed genes. Cells were classified into five shared categories, epithelial cells, fibroblasts, endothelial cells, macrophages, and lymphocytes, using discrete marker-based classification for PCF and probabilistic assignments for MERSCOPE (Fig. 4b, Fig. S6).

We applied SAME to integrate the PCF and MERSCOPE modalities, using the optimal hyperparameters found in the previous section. SAME’s capacity to integrate discrete (protein) and probabilistic (mRNA) cell classifications enabled accurate spatial correspondence even in the presence of dropout and resolution differences, with 78% of cells successfully matched between modalities. This robustness was maintained even when accounting for the inherent noise in RNA expression measurements, with PCF cell types appearing within the top 2 and top 3 MERSCOPE predictions in 85.1% and 89.9% of cases, respectively (Fig. S7). A merged expression heatmap (Fig. 4c) demonstrates strong spatial concordance between protein and mRNA markers within cell types. Epithelial markers such as PanCK aligned precisely with the keratins KRT6A and KRT4, while other lineage-defining markers showed coherent organization across spatially aligned neighborhoods. This validates the effectiveness of SAME in preserving spatial structure while enabling biologically meaningful multimodal integration.

Leveraging the spatially aligned dataset, we focused on a defined lymphoid niche within the dorsal tongue epithelium (Fig. 4d–f). Protein markers allowed reliable identification of T cells, and their spatial coordinates were used to distinguish epithelial-resident T cells from the epithelial and also from deeper non-epithelial counterparts.

Differential expression analysis revealed that epithelial-resident T cells upregulated IL1RN and NOTCH3, consistent with roles in local inflammation control and tissue-specific adaptation. Notably, these cells also showed elevated expression of IGFBP2, IGFBP3, and IGFBP6, suggesting a transcriptional program oriented toward metabolic regulation, and modulation of growth factor signaling. These features are consistent with a model where epithelial-resident T cells adopt a non-proliferative state to maintain barrier function under persistent environmental exposure. In contrast, T cells in the parenchyma showed elevated levels of IL10RA and IGFBP4, indicating a different strategy for long-term suppression of immune activation. IL10RA confers responsiveness to IL-10, a cytokine central to immune regulation and tolerance, while IGFBP4 has been associated with cell quiescence and negative regulation of growth and survival pathways. Rather than responding to acute environmental stimuli, these cells appear tuned to maintain tissue homeostasis.

Together, these findings reveal that while both subsets of T cells contribute to tissue protection, they do so through *distinct mechanistic pathways shaped by their spatial niche*: epithelial-resident T cells actively suppress barrier-localized inflammation and improve survival under environmental stress, whereas parenchymal T cells adopt a quiescent, regulation-oriented phenotype to maintain long-term tissue equilibrium. Additional investigations could reveal how inflammation or injury result in differing responses within the two spatial niches.

These insights were critically dependent on cross-modal integration. Classification based solely on MERSCOPE data yielded low-confidence lymphoid assignments due to limited transcript detection, which would have obscured both the presence and functional heterogeneity of tissue-resident T cells. By anchoring cell identity to high-confidence protein-based classification and enriching this with transcriptional information, SAME enabled the detection of cryptic immune populations and their spatially determined gene programs.

### Discovering niche-dependent T cell states in lung adenocarcinoma

Building on our results in healthy tissue, we next examined whether SAME-mediated multimodal integration could uncover niche aligned variation in T cell phenotypes within diseased, highly heterogeneous tumor environments. Unlike normal epithelial sites, solid tumors such as lung adenocarcinoma present a profoundly disordered landscape, characterized by disrupted barriers, architectural disarray, and patchy immune infiltration [28]. Such complexity poses substantial challenges for cell-type deconvolution and functional annotation when using RNA-based profiling alone, particularly in regions densely infiltrated by immune or cancer cell populations.

To address these challenges, we applied SAME to pairwise-align protein and transcriptomic data from two consecutive sections of lung adenocarcinoma, stained with highly multiplexed antibody panels (PhenoCycler-Fusion) and spatial mRNA probes (Xenium) respectively [29]. After stringent segmentation and quality control, both modalities yielded comparable cell counts (94,442 for PCF and 99,827 for Xenium), and application of the multimodal phenotyping pipeline identified six principal cell classes, including epithelial (tumor), mesenchymal, macrophage, B cell, and T cell populations (Fig. 5a, Fig. S8–S9). As observed in healthy tissue, T cells were substantially underrepresented in the RNA dataset (3,031 cells) relative to protein (8,052 cells), again emphasizing the necessity of cross-modal alignment for accurate cell-type recovery.

**Figure 5.**
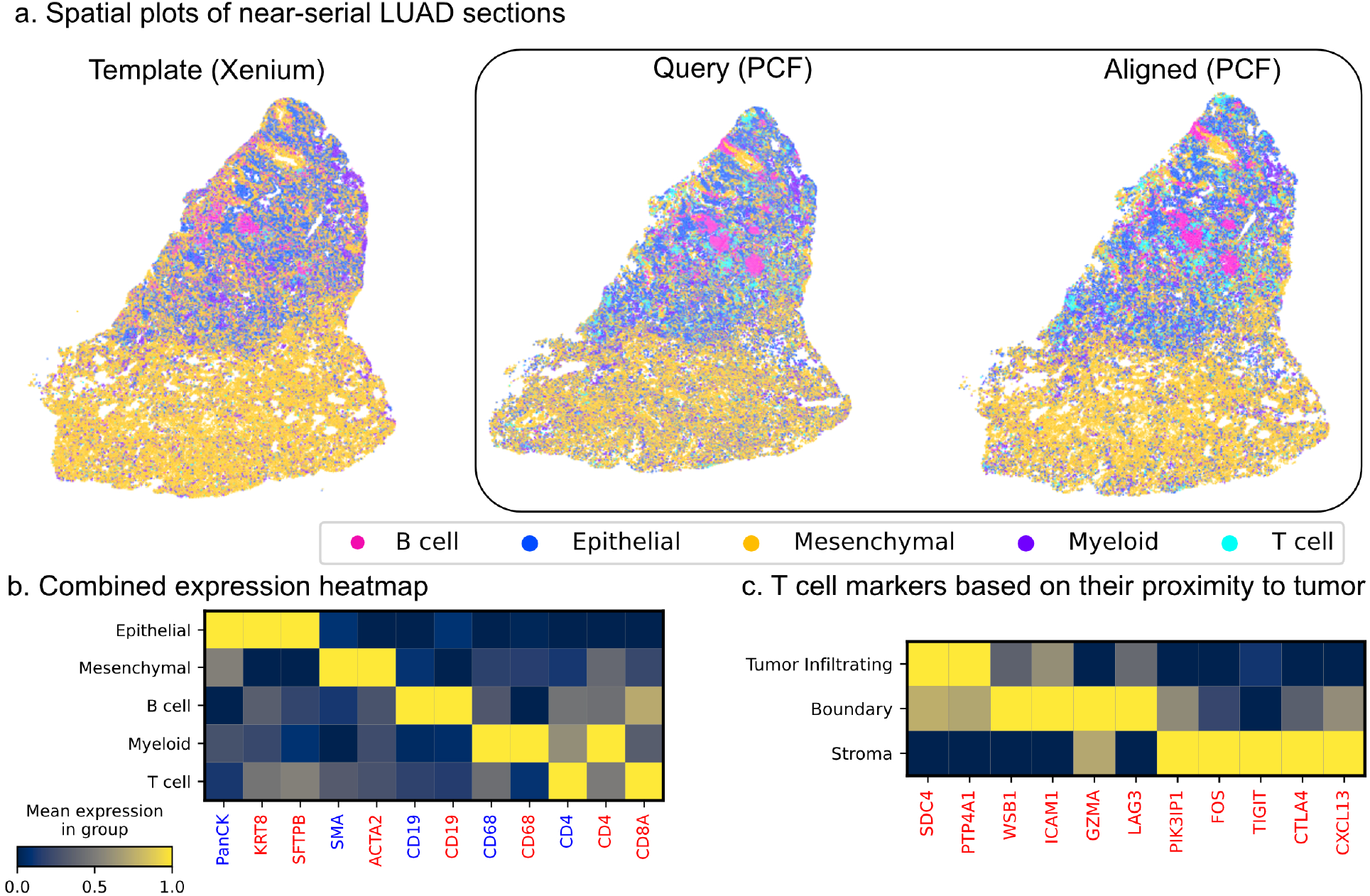
Integrated spatial mapping of protein and RNA in lung adenocarcinoma. a. Spatial cell-type plots for two near-serial lung adenocarcinoma sections: Template (Xenium, RNA), Query (PCF, protein), and PCF after SAME alignment, showing major cell types. b. Heatmap of combined SAME expression profiles with protein markers (blue labels) and transferred mRNA (red labels) across major cell types. c. Heatmap of T cell marker expression stratified by proximity to tumor cells, highlighting distinct transcriptional programs in tumor-infiltrating, boundary, and stromal T cell compartments.

SAME alignment preserved spatial integrity and demonstrated robust concordance between protein and mRNA markers for key cell types (Fig. 5b, Fig. S10). This enabled us to focus on the spatial distribution and specialization of T cells throughout the tumor microenvironment. By mapping T cell positions relative to neoplastic regions (Methods), we distinguished three spatially defined subgroups: intratumoral T cells (within 15 *µ*m of tumor cells), peritumoral/boundary T cells (15–30 *µ*m), and stromal T cells (> 30 *µ*m from tumor cells).

Differential gene expression analyses among these groups revealed marked microenvironmental shaping of T cell programs (Fig. 5c). Intratumoral T cells exhibited pronounced enrichment of SDC4 and PTP4A1—genes linked to blunted T cell activation and regulatory processes, suggesting a profile consistent with chronic stimulation and locally imposed restraint. T cells at the tumor boundary showed upregulation of WSB1, ICAM1, GZMA, and LAG3, together indicating dynamic engagement with their environment: elevated adhesion (ICAM1), cytotoxic activity (GZMA), and early induction of exhaustion markers such as LAG3. This hints at a transitional phenotype—simultaneously poised for effector function yet increasingly constrained by suppressive signals.

In contrast, stromal T cells were marked by high expression of canonical suppressive or regulatory genes, including PIK3IP1, FOS, TIGIT, CTLA4, and CXCL13. This gene signature suggests attenuation of T cell activation and support for immune quiescence or tertiary lymphoid structure formation, in line with functional detuning under sustained exposure to suppressive cues within the stromal compartment.

Together, these findings highlight the power of integrated spatial profiling in revealing how the tumor microenvironment orchestrates spatially-stereotyped immune cell functional states. By leveraging robust cell-type annotation from protein data and resolving functional heterogeneity at transcriptomic depth, SAME uncovers context-dependent T cell adaptation that would be otherwise masked in single-modality analyses. These results underscore the importance of spatial context for immune specialization in cancer, and establish a generalizable platform for microenvironmental dissection in complex pathological tissues.

### Elucidating metabolic mechanisms of TAM-mediated immune evasion in lung adenocarcinoma

Metabolic reprogramming within tumor microenvironments drives cancer progression and immune evasion [30, 31], yet characterizing these processes at cellular resolution remains challenging. Spatially resolved metabolomics techniques like MALDI-MSI can capture metabolic landscapes across tissues [32, 33, 34], but determining which cell types produce specific metabolites is difficult without complementary cell-type information. Conversely, protein-based spatial profiling provides robust cell-type identification but lacks insight into metabolic activity. These limitations have hindered comprehensive understanding of metabolic cross-talk between specific cell populations within tumor niches.

To this end, we assayed two near-serial sections of lung adenocarcinoma with spatial protein and metabolic profiling to expand SAME’s integration capabilities across these modalities (Fig. 6a–b). Spatial protein profiling using multiplexed immunofluorescence (PCF) clearly identified major cell populations based on canonical markers (Fig. 6b). However, since metabolic markers alone lack the specificity needed for reliable cell-type identification, we extracted morphological features from H&E images using a pathology foundation model [19], then assigned cell types based on the closest PCF-defined populations after an initial affine transform-based alignment (Methods, Fig. S11.). We then applied SAME to estimate comprehensive, cell-type specific molecular signatures. Fig. 6c illustrates these integrated signatures, showcasing distinct protein and metabolite levels characteristic of each major cell types identified in PCF (Fig. 6d). This demonstrates SAME’s ability to align and assign metabolic information to proteomically-defined cell populations, thereby enriching the characterization of each cell type.

**Figure 6.**
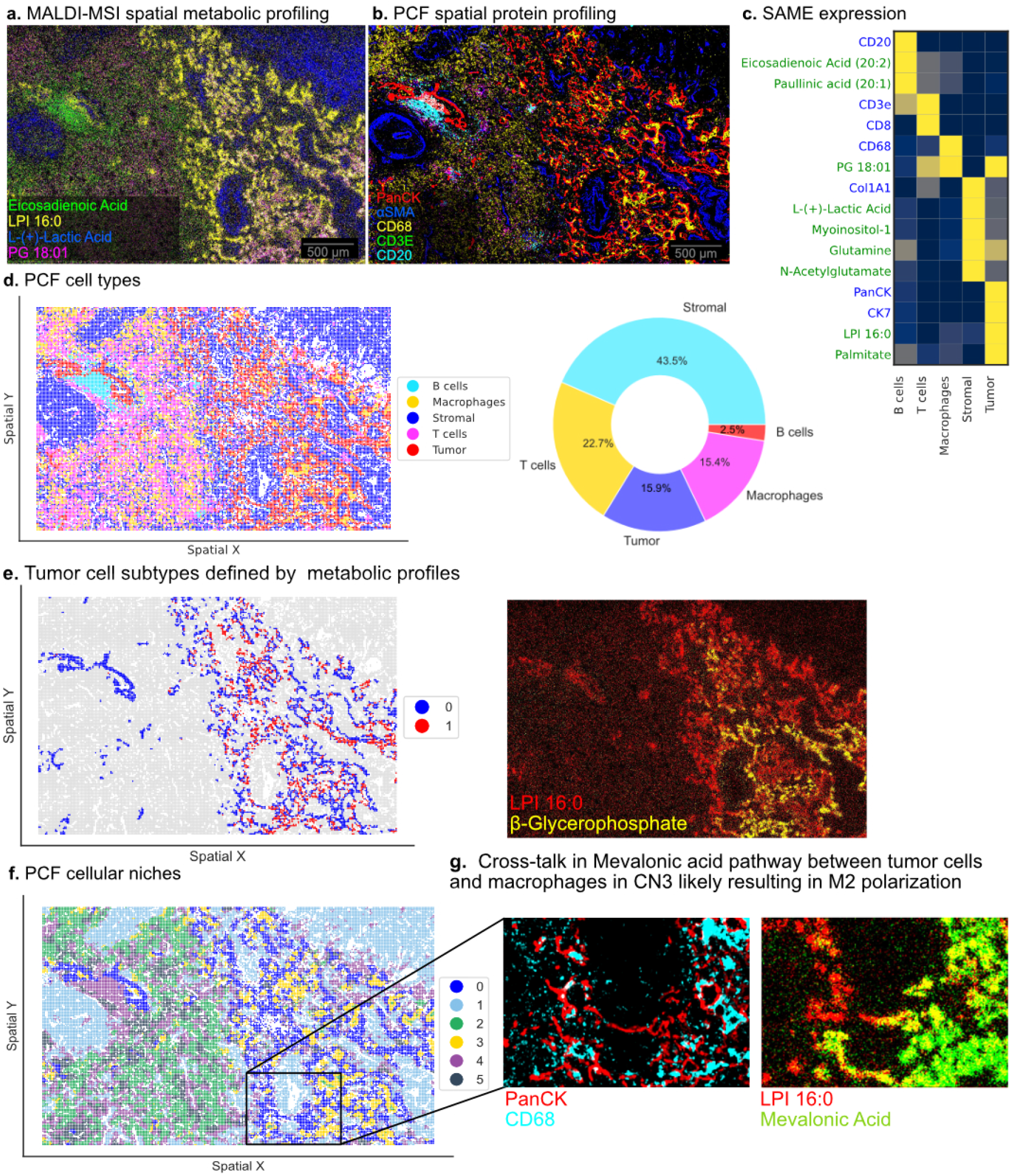
SAME integration of spatial protein and spatial metabolome profiling on two near serial sections from lung adenocarcinoma. a. Spatial metabolome profiling of differentially enriched candidate markers showing broad cell-type specific patterns. b. Spatial protein profiling using multiplexed immunofluorescence imaging using PCF showing five markers PanCK (red), aSMA (blue), CD68 (yellow), CD3E (green), CD20 (cyan) representing Tumor, stroma, macrophage, T, and B cell populations. Pie-chart shows the cell-type proportions (n=39,855). c. Heatmap showing cell-type specific protein (blue) and metabolite (green) expression in the major cell types after SAME-based integration. d. Spatial scatter plot showing cell types identified using PCF. e. Unsupervised clustering of metabolic profiles of tumor cell population identifies two major subtypes (left) and two markers indicative of differences in metabolite levels (right). f. Cellular niche analysis showing six major cell niches. g. Zoomed-in region showing cells from CN3 comprising primarily of tumor cells and macrophages and up-regulated Mevalonic acid metabolite expression indicating potential cross-talk.

Beyond broad cell-type characterization, the integrated data allowed for deeper investigation of intra-tumoral heterogeneity. Unsupervised clustering of SAME-aligned metabolic profiles from Pan-cytokeratin positive tumor cells revealed at least two major metabolic subtypes (Fig. 6e, left). Differential metabolite analysis between these tumor subtypes highlighted specific metabolites, such as distinct levels of *β*-Glycerophosphate and Lysophosphatidylinositol (LPI) 16:0 (Fig. 6e, right). Alterations in *β*-Glycerophosphate may indicate differences in phospholipid metabolism, crucial for cell membrane biogenesis and remodeling during proliferation, or reflect variances in cellular signaling pathways where phosphorylated intermediates play key roles [35]. Concurrently, differing LPI 16:0 levels within lung adenocarcinoma tissue suggest metabolic heterogeneity among tumor cells; elevated LPI 16:0 levels in tumor cells may contribute to enhanced proliferative and survival capacities, potentially as an adaptive response to microenvironmental stressors [36, 37].

Furthermore, the integrated multimodal data revealed a diverse set of cellular niches within the tumor microenvironment. We identified six major cellular niches based on the co-occurrence and spatial relationships of different cell types (Fig. 6f). Notably, Cellular Niche 3 (CN3) is primarily composed of tumor cells and macrophages (Fig. 6g, zoomed-in region). Within this tumor-macrophage rich niche, we observed upregulated expression of Mevalonic acid. The mevalonate pathway, for which Mevalonic acid is a key intermediate, is critical for synthesizing cholesterol and govern cell proliferation, survival, and motility. Upregulation of the mevalonate pathway is a common feature in many cancers, including lung adenocarcinoma, where it fuels tumor growth and aggressiveness [38, 39]. Its heightened activity specifically within the CN3 tumor-macrophage hotspot suggests a site of intense anabolic activity and signaling. This localized metabolic alteration points towards a synergistic metabolic relationship or cross-talk, where tumor cells and macrophages might mutually support or induce this pathway [40]. For instance, macrophages can be “educated” by tumor cells to adopt a pro-tumor phenotype [41, 42, 43] and, in turn, might secrete factors that stimulate tumor cell proliferation and mevalonate pathway activity, or even supply precursors [44, 40].

## Discussion

Spatial multiomics integration aims to leverage complementary modality strengths while accommodating their conflicting processing protocols. However, different modalities often require conflicting optimal protocols. For example, protein detection benefits from specific antibody adsorption conditions that may compromise RNA integrity, while metabolomics typically demands fresh-frozen processing unsuitable for other assays [1, 45]. Serial sectioning thus enables experimental designs impossible with same-section approaches. Even when assays are repeated on the same tissue section, repeated handling and imaging can introduce spatial artifacts [46]. SAME enables researchers to process each section optimally for its intended modality, then computationally recover spatial relationships. Importantly, SAME’s flexible formulation supports integration of diverse spatial modalities, including transcriptomic, proteomic, metabolic, and even histological image-based inputs, as demonstrated through our integration of PCF protein data with MERSCOPE transcriptomics, Xenium RNA profiling, and MALDI-MSI metabolomics across different tissue types.

Current spatial alignment methods fail when confronted with the topological violations common in real tissue processing. PASTE’s linear transformation assumptions cannot accommodate local distortions, limiting its application to sections with minimal deformation [9, 10]. GPSA [11] and STalign [13] enforce topology-preserving constraints that, while mathematically elegant for smooth deformations, cannot handle tissue tears, holes, or dropped regions. MIIT attempts multiple serial section integration based on histology images, but it is still limited to diffeomorphic registration, so it cannot handle tissue tears or missing regions, forcing users to exclude problematic areas and lose spatial coverage [47]. In contrast, SLAT’s graph neural network approach permits arbitrary topological changes through adversarial matching but provides no mechanism to detect or control tissue topology violations [48]. Space-tearing transforms in SAME address this pressing need for a principled approach to controlled topological flexibility in spatial integration. Our triangle orientation constraint provides a geometric measure of local topological change, while integer programming balances cell-type correspondence against spatial coherence through tunable penalties. SAME thus generalizes existing frameworks—its phenotype-correspondence approach applies to both multimodal and unimodal settings, with the extent of space-tearing configurable from zero (i.e., a topology-preserving transform) to high, based on the data requirements.

SAME’s abilities are critical for exploring biological variables that often require multiple modalities. For example, certain cell states are defined by the localization pattern (nucleo-cytoplasmic shuttling), post-translational modifications, or polarization (eg: apical membrane or fillipodia) of some proteins, which can only be inferred from protein based assays. Whereas, characterizing the expression of non-coding transcripts (eg: long non-coding RNA, miRNA) require RNA based assays. Additionally, most high-resolution spatial transcriptome profiling methods lack depth, making them not sufficient to define certain cellular phenotypes such as activation state of a cell. Our finding that IL1RN and IL10RA expressing lymphocytes in intraepithelial versus sub-epithelial microniches, respectively, was only possible by integrating protein-based cell typing (CD4 protein expression) with high-resolution transcriptional profiling (IL10RN and IL10RA). Further, with the advent of MALDI-MS imaging it is now possible to profile small molecule based metabolites, lipids, and different types of glycocalyx, which require other modalities for cell segmentation and annotation. Indeed, as we demonstrated here SAME unlocks the power of emerging spatial metabolomics by enabling integration with established protein profiling. Our lung adenocarcinoma study, integrated spatial metabolites and multiplexed protein profiling, uncovered localized upregulation of mevalonic acid specifically within tumor-macrophage niches. This suggests targeted metabolic crosstalk invisible to bulk approaches and demonstrates how multimodal integration can uncover regulatory mechanisms operating at cellular neighborhood scales.

The power of SAME derives also from a careful incorporation of additional data sources and by exploiting computational advances such as foundation models. Our phenotype matching approach makes SAME modality-agnostic, enabling expansion to future technologies. Moreover, SAME transforms the challenge of noisy measurements into an advantage–rather than requiring definitive cell-type assignments, SAME naturally incorporates probabilistic phenotype scores into its optimization framework, yielding uncertainty-aware correspondences that reflect the confidence in both molecular measurements and spatial proximity. Introduced for metabolite integration, our use of H&E foundation models [19, 20] to extract phenotypic features from morphology offers a general strategy for modalities lacking direct cell-type information. Leveraging the extensive research in medical image registration [14, 49], we exploit broadly-available readouts like DAPI and H&E to compute a starting alignment.

SAME balances computational complexity with practical performance through strategic algorithmic design. The constraint generation phase has 𝒪 (*nk*^3^) complexity—where *n* represents the number of cells in the query dataset and *k* represents the maximum number of potential matches considered per cell in the template dataset. The core 0-1 ILP optimization belongs to the NP-hard complexity class, theoretically requiring exponential time in the worst case (Note S1). However, modern ILP solvers like Gurobi employ sophisticated branch-and-bound algorithms that rapidly converge to near-optimal solutions in practice. We further accelerate computation through a patch-based approach that divides tissue sections into overlapping regions, solving local subproblems before global integration. For the ISS heart dataset with a few thousand cells per section, SAME completes alignment in under 11 hours on standard computational infrastructure in the most restrictive scenario, where we desire space-tearing transforms to less than 5% (Fig. S3b). Runtime can be reduced further by relaxing constraints. For example, permitting approximately 10% space-tearing transforms reduces computation time by half to around six hours (Fig. S3b). Future implementations could leverage approximate optimization methods or GPU acceleration to further improve scalability.for datasets exceeding current computational limits.

SAME is modality agnostic, enabling experimental designs where each modality can be processed under their optimal conditions. This capability becomes increasingly important as spatial omics technologies mature and researchers demand comprehensive molecular characterization of complex tissues. The framework’s modality-agnostic design, positions SAME to integrate emerging spatial technologies including spatial ATAC-seq [50] and spatial chromatin imaging [51]. SAME’s ‘s controlled topological flexibility provides a foundation for comprehensive molecular characterization of complex tissues, from developmental atlases capturing rapid morphogenetic changes to disease progression studies requiring robust alignment across heterogeneous pathological architectures.

## Methods

### SAME Problem Formulation

Given a template slide *T* = {*t*_1_, *t*_2_, …, *t* _|_*T*_|_} and a query slide *Q* = {*q*_1_, *q*_2_, …, *q*_|_ *Q*_|_}, where each cell has spatial coordinates (*x*_*i*_, *y*_*i*_) and phenotype probabilities, we formulate the alignment as a constrained assignment problem. Here, (*x*_*i*_, *y*_*i*_) denote the spatial coordinates of cell *i*, and *u*_*ij*_ are binary assignment variables. The optimization seeks an assignment matrix *U* that minimizes:

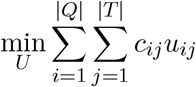

subject to:

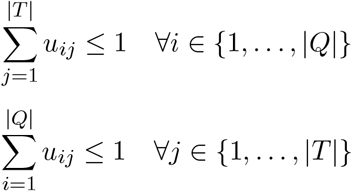

where *u*_*ij*_ ∈ {0, 1} indicates whether query cell *q*_*i*_ matches template cell *t*_*j*_. The cost function *c*_*ij*_ combines phenotypic correspondence and spatial proximity:

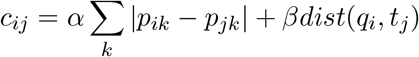

where *p*_*ik*_ represents the probability that the cell *i* belongs to the phenotype 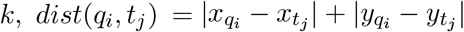 is the L1-distance between the spatial coordinates of the query cell *q*_*i*_ and the template cell *t*_*j*_, and *α, β* weight the relative importance of phenotypic similarity versus spatial distance.

### Space-Tearing Through Triangle Orientation Constraints

The key innovation in SAME lies in monitoring and controlling topological violations through triangle orientation changes. We construct a Delaunay triangulation *D* over query cells, where each triangle *τ* = (*q*_*a*_, *q*_*b*_, *q*_*c*_) ∈ *D* captures local geometric relationships. The signed area of triangle *τ* is computed as:

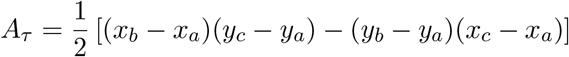

The sign of *A*_*τ*_ indicates whether vertices are ordered clockwise (*A*_*τ*_ *<* 0) or counterclockwise (*A*_*τ*_ *>* 0). A sign flip after mapping indicates a topological violation—a space tear.

For each triangle *τ* ∈ *D* and each possible mapping to template cells 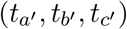, we introduce constraints:

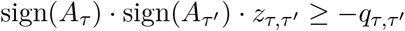

where:

- 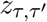 is a binary variable that equals 1 if and only if all three vertices of *τ* map to *τ*^*′*^ (see below),
- 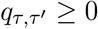 is a penalty variable allowing orientation violations,
- 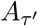 is the signed area of the mapped triangle in template space.

The product of binary variables 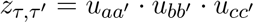 is linearized as follows:

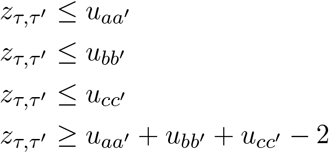

These constraints ensure 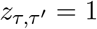 if and only if all three vertices of *τ* are simultaneously mapped to the corresponding vertices of *τ*^*′*^.

### Complete Objective Function

The full optimization balances phenotypic correspondence against geometric distortion:

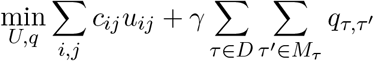

where *γ* controls the penalty for topological violations, and *M*_*τ*_ represents valid template triangles for *τ*.

### Computational Efficiency Strategies

We implement the following strategies to considerably reduce the search space:

1. **Candidate Pair Restriction:** Rather than considering all |*Q*| *×* |*T*| possible matches, we restrict candidates using a *k*-nearest neighbor graph. For each query cell *q*_*i*_, we identify the *k* nearest template cells within radius *r*.This reduces the search space from 𝒪 (|*Q*| |*T*|) to 𝒪 (*k*| *Q*|), where *k* is typically 8 cells.
2. **Delaunay Triangle Pruning:** We retain only on triangles obtained using Delaunay triangulation with maximum edge length below threshold *ρ*, focusing on local spatial relationships. This filtering step ensures that only non-degenerate, local triangles contribute to orientation-based constraints in the optimization. Further, in all our experimentally-derived spatial datasets, we exclude triangles if the triangle vertices are composed entirely of the same cell type, as they provide limited information for space-tearing violations.
3. **Soft Constraint Formulation:** All spatial constraints are implemented as soft constraints through penalty variables. This ensures feasibility even when perfect topology preservation is impossible due to biological variation or technical artifacts. Hard constraints can be recovered by setting penalty coefficients arbitrarily high.
4. **Sliding Window Decomposition:** For large tissue sections, we offer a sliding window approach. The tissue is partitioned into overlapping windows of size *w × w* pixels with overlap *δ*:

- Each window is optimized independently using the full SAME formulation
- Matches are retained only from the central (*w* − 2*δ*) *×* (*w* − 2*δ*) region
- Windows with fewer than five cells are merged with adjacent windows
- Results are aggregated into a global solution. Further, this decomposition enables parallel processing while maintaining spatial continuity across window boundaries.

### Implementation Details

We solve the integer linear program using Gurobi v12.0 with a MIP gap tolerance of 0.5% and a time limit of 7200 seconds per window. The default penalty coefficient *γ* is set to 10 but can be tuned based on data quality and the desired trade-off between phenotypic correspondence and spatial coherence. To focus on local spatial relationships, we set the triangle edge threshold *ρ* to five times the median cell-cell distance in the dataset. The optimization leverages Gurobi’s branch-and-cut algorithm with warm starts from the initial image-based alignment. For typical window sizes containing 100-500 cells, optimal or near-optimal solutions are found within minutes, enabling efficient processing of large tissue sections.

### Datasets

We evaluated our approach on several synthetic and datasets from spatial genomic, proteomic, and metabolomic datasets described below.

### Synthetic warps

We generated two different types of synthetic warps starting with a regular 12×12 grid of cells with each alternate cell being assigned one of the two cell types. The two warps are generated as follows: To obtain the distorted slice, we applied two types of spatial distortion:

1. **Diffeomorphic Warp**: We obtained a diffeomorphic warp using a smooth, spatially correlated distortion using a Gaussian Process with an RBF kernel (length scale=1, variance=0.1). This simulates coherent tissue deformation that might occur during sample preparation. The distortion can be modeled as:

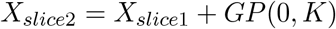

•where *K* = 0.1 · *RBF* (*scale* = 1) is the kernel function applied to the original coordinates.
2. **Space-tearing Warp**: We started with the diffeomorphic warp as above and applied additional random displacement of up to 0.5 units in magnitude in the bottom right corner of the synthetic tissue slice. This simulates local distortions and space-tearing distortions that might affect individual cells or regions differently:

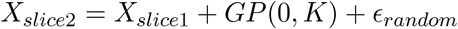

• where *ϵ*_*random*_ is applied only to randomly selected points with probability 0.5.

### Near-serial embryonic heart dataset

We analyzed spatial transcriptomic data from two near-serial sections of embryonic heart tissue at 6.5 post-conception weeks (pcw), profiled using single-molecule fluorescence in situ hybridization (smFISH) [21]. The authors provided detailed cell phenotype annotations based on RNA markers. To establish template and query designations, we selected the section with minimal tissue distortions and folds as the template, while the section with more artifacts served as the query. Initial spatial alignment between sections was performed using Valis [16] to register DAPI nuclear staining images acquired during in situ sequencing (ISS). This VALIS-aligned query sample served as the ‘Initial’ starting point for all alignment algorithms evaluated in this study.

### Sample Preparation and Data Acquisition for Dorsal Tongue

Multiplexed immunofluorescence (PhenoCycler-Fusion) was conducted using the PhenoCycler-Fusion 2.0 platform (Akoya Biosciences). FFPE tissue sections were first deparaffinized through a graded ethanol series (100% - 30%). Antigen retrieval was carried out in an AR9 buffer (EDTA-based, Akoya) using a low-pressure cooker for 15 minutes. After cooling the slides for 1 hour at room temperature, they were briefly rehydrated in 100% ethanol for 2 minutes and then incubated for 20 minutes in a staining buffer. Antibody cocktails were prepared following the PhenoCycler-Fusion user guide, combining four blocking agents J, G, N and S nuclease-free water, and staining buffer. Primary antibodies were diluted at 1:200 in this cocktail. A combined mix of all primary antibodies was applied to the slides, which were incubated overnight at 4°C in a humidified chamber (Sigma-Aldrich).

Post-incubation, slides were briefly washed in a staining buffer (2 minutes) and fixed in 10% paraformaldehyde (PFA) diluted in a staining buffer for 10 minutes. This was followed by three washes in 1X PBS (2 minutes each). The sections were then treated with ice-cold methanol for 5 minutes. A final fixative solution (FFS), as specified in the PhenoCycler protocol, was applied to the slides for 20 minutes at room temperature. After an additional PBS wash, slides were transferred to the FCAD module for flow cell assembly. A brief high-pressure application (30 seconds) secured the flow cell on the slide.

Following mounting, slides were incubated in a PCF buffer for 10 minutes and loaded onto the PhenoCycler-Fusion instrument with 41 antibodies (Tab. S1). Reporter solutions were prepared fresh for each cycle using stock reporter mix, nuclease-free water, 10X PCF buffer, PCF assay reagent, and DAPI to reach a final 1:1000 DAPI dilution. Two slides were processed per run. Each reporter was diluted 1:50 in the final reaction, and 250 *µ*L per reporter were loaded into a 96-well plate, which was sealed with aluminum foil (Akoya). Manual mapping of the scanning region was performed using brightfield mode on the PhenoImager. DMSO concentrations (low and high) were prepared according to Index B specifications in the PhenoCycler-Fusion manual.

Multiplexed error-robust fluorescence in situ hybridization (MERFISH) was carried out using the MERSCOPE system (Vizgen). To confirm assay performance and signal reliability, quality control was performed on representative specimens from each oral niche utilizing the MERSCOPE Verification Kit (Vizgen), following the manufacturer’s guidelines (91600004 Rev D) and pipeline previously described by Matuck et. al [25].

For cell type annotation, we employed different strategies for each modality because protein markers provide more specific and reliable cell type identification compared to the inherently noisier transcriptomic readouts. For the MERSCOPE-based slide, we trained a logistic regression classifier using the reference scRNA-seq dataset for the dorsal tongue obtained from matching dorsal tongue tissue [25] and classified cells into the five broad categories. We retained the predicted probabilities for each cell type for downstream analysis. For the PCF data, we performed DAPI-based nuclear segmentation and intensity quantification using QuPath’s cell detection plugin [49]. After normalizing the data with Scanpy [52] (using log1p and scale transformations), we performed Leiden clustering and manually annotated the resulting clusters with the same five cell type categories used for the MERSCOPE data, ensuring consistency for SAME-based alignment.

### Near-serial Lung Adenocarcinoma dataset for Protein-RNA Alignment

We analyzed two near-serial sections from lung adenocarcinoma specimen LUAD No. 33, a minimally invasive adenocarcinoma (MIA) classified as Noguchi Type C [29]. The first section was profiled using Xenium spatial transcriptomics with a 302-gene panel, while the adjacent section underwent PhenoCycler-Fusion (PCF) multiplexed protein profiling with 35 antibodies.

Similar to the ‘Dorsal Tongue’ tissue analysis, we employed different strategies for each modality. For the Xenium sample, we trained a logistic regression classifier using the reference scRNA-seq dataset provided by Takano et al. [29, 53], focusing on five broad cell type categories. We retained the predicted probabilities for each cell type for downstream analysis. The PCF data underwent the same processing pipeline as described in the ‘Dorsal Tongue’ section, yielding five consistent cell type categories for SAME alignment.

### Data Acquisition for Lung Adenocarcinoma sample for Protein-Metabolomic Alignment

#### Tissue Collection and Sectioning

Human lung tissue samples were obtained from surgical resections of lung adenocarcinoma patients. The study protocol was approved by the ethics committee (“Ethik Kommission am Fachbereich Humanmedizin der Justus Liebig Universität Giessen”) in accordance with national regulations and Good Clinical Practice/International Conference on Harmonisation (GCP/ICH) guidelines. Written informed consent was obtained from all participants (ethical reference number: AZ 58/15).

Tissue specimens were immediately snap-frozen in liquid nitrogen and stored at -80°C until further processing. The samples were embedded in 8–10% gelatin before sectioning. Cryosections (10–12 *µ*m thick) were prepared at -20°C using a Thermo CryoStar NX50 cryostat and thaw-mounted onto indium tin oxide (ITO)-coated conductive glass slides (Bruker Daltonics). Sections were then desiccated under vacuum for at least 30 minutes to remove residual moisture and improve matrix adherence.

#### PCF Sample Preparation and Data Acquisition

Serial sections (5*µ*m thickness) of fresh frozen (FF) tissue specimens were obtained using a cryostat (Thermo Scientific, CryoStar NX50) and mounted centrally on Polysine coated glass slides (Epredia, Cat#J2800AMNZ). Prepared slides were stored at -80°C until further use.

Prior to antibody staining, frozen samples were removed from storage and equilibrated at room temperature (RT) for 2 minutes on a bed of Drierite desiccant. Samples were then fixed in acetone (Sigma-Aldrich, Cat# 34850) for 10 minutes at RT, followed by a 2-minute air-drying step at RT. Subsequently, the samples were transferred into Hydration Buffer in preparation for the application of Parhelia Cover Pads (Parhelia Biosciences, Cat# PN40221). For antibody staining, the antibody cocktail (containing 30 PhenoCycler antibodies, Tab. S2) was prepared with optimal dilutions of each antibody in a Staining Buffer containing G, S, J and N blockers. Tissue staining was performed using the Parhelia Spatial Station (PSS), with the PhenoCycler-Fusion staining protocol executed in a fully automated manner on the PSS platform with staining reagents from Akoya Biosciences (Akoya Biosciences, part of kit-Cat#7000017).

After staining was completed, samples were removed from PSS and Flow Cells (Akoya Biosciences, part of kit-Cat#7000017) were attached to each sample. A PhenoCycler-Fusion reporter plate was prepared, and a PhenoCycler-Fusion run was initiated. Images were acquired and processed, resulting in the generation of the final .qptiff file.

#### MALDI-MSI Sample Preparation and Data Acquisition

Matrix coating was performed using 1,5-diaminonaphthalene (DAN), prepared at a concentration of 10 mg/mL in a 70:30 (v/v) acetonitrile:water solution. The matrix was uniformly applied using an HTX TM-Sprayer (HTX Technologies, USA) under the following conditions: 8 passes, flow rate of 70*µ*L/min, nozzle temperature of 70°C, and track spacing of 2mm [54].

Mass spectrometry imaging was performed using a Bruker rapifleX MALDI-TOF mass spectrometer operating in negative ion mode. Spectra were acquired over a mass-to-charge (m/z) range of 50–650, with a spatial resolution of 5*µ*m per pixel, enabling high-resolution molecular imaging. Each pixel spectrum was generated by accumulating 15 laser shots. External calibration was conducted prior to acquisition using red phosphorus, ensuring accurate m/z assignment throughout the dataset.

Raw MALDI imaging data were imported into SCiLS Lab 2025b Core software (Bruker Daltonics, Bremen, Germany) for preprocessing. Baseline correction and Root Mean Square (RMS) normalization were uniformly applied across all datasets to minimize technical variability and enable robust inter-sample comparisons. Unsupervised segmentation was carried out using bisecting *k*-means clustering in SCiLS Lab to identify spatially distinct molecular regions. These were aligned with corresponding hematoxylin and eosin (H&E)-stained images obtained from the same tissue sections [55]. For metabolite annotation, m/z features were cross-referenced using the MetaboScape 2025 platform (Bruker Daltonics), employing a combination of in-house and public spectral libraries. The results were first exported in .mca format to SCiLS Lab for peak width refinement and ion image quality assessment, then exported in OME-TIFF format for downstream spatial analyses.

#### Data pre-processing

Integrating MALDI-MSI metabolomics data with other spatial modalities presents a fundamental challenge: metabolite intensities cannot be directly translated to cell types. We developed a preprocessing pipeline that leverages ubiquitous H&E staining to extract spatial features that serve as phenotype proxies. We applied H-optimus-0, a foundation model for histology trained on over 500,000 H&E stained whole slide images. Each 224×224 pixel patch (0.5 *µ*m resolution) in the H&E image was converted to a 14×14×1536 feature tensor, which we then rescaled to match the MALDI-MSI spatial resolution (Fig. S11a-b).

We processed these patch-level features as spot-based spatial data: principal component analysis reduced dimensionality while preserving 67.3% of morphological variance, followed by unsupervised clustering that identified 13 spatially coherent regions (Fig. S11c). To validate that these morphological clusters captured biologically relevant cell populations, we performed affine alignment with a serial section containing immunofluorescence data. The resulting cluster-to-cell type mapping revealed distinct associations: clusters 0 and 8 corresponded to T cells and macrophages, cluster 6 to lymphoid aggregates, clusters 3 and 10 to tumor epithelium, and remaining clusters to various stromal components (Fig. S11d). For direct comparison, we downsampled the PCF serial section to match the MALDI-MSI resolution and performed unsupervised clustering, which recovered similar cell type distributions (Fig. S11e). This morphology-based phenotyping approach enabled SAME to successfully align metabolomics data with spatial transcriptomics, bridging modalities that lack direct molecular correspondence.

## Code and Data availability

The implementation of SAME is available at: https://github.com/rohitsinghlab/SAME

## Acknowledgements

A.P. and R.S. acknowledge the support of the Whitehead Scholarship from the Duke University School of Medicine, as well as the Chan Zuckerberg Initiative. P.R.T. is partly supported by Chan Zuckerberg Initiative DAF award 2022-237918.

## Supplementary Data for ‘SAME: Topology-flexible transforms enable robust integration of multimodal spatial omics’

### Note S1: Computational Complexity of SAME

The space complexity of SAME’s setup phase is dominated by the spatial constraint generation step, which requires 𝒪 (*nk*^3^) memory to store all constraint variables and constraint mappings, where *n* represents the number of cells in the query dataset (*Q*) and *k* is the number of nearest neighbors in the template dataset (*T*) considered per cell. This cubic dependence on *k* arises from the need to enumerate all possible matching combinations for the three vertices of each Delaunay triangle, with each vertex having up to *k* candidate matches in the reference dataset. The constraint generation process creates auxiliary binary variables for each valid triangle-to-triangle correspondence, along with associated penalty variables for spatial violations, resulting in the storage of approximately 𝒪 (*nk*^3^) variables and their associated constraint coefficients.

**Figure S1.**
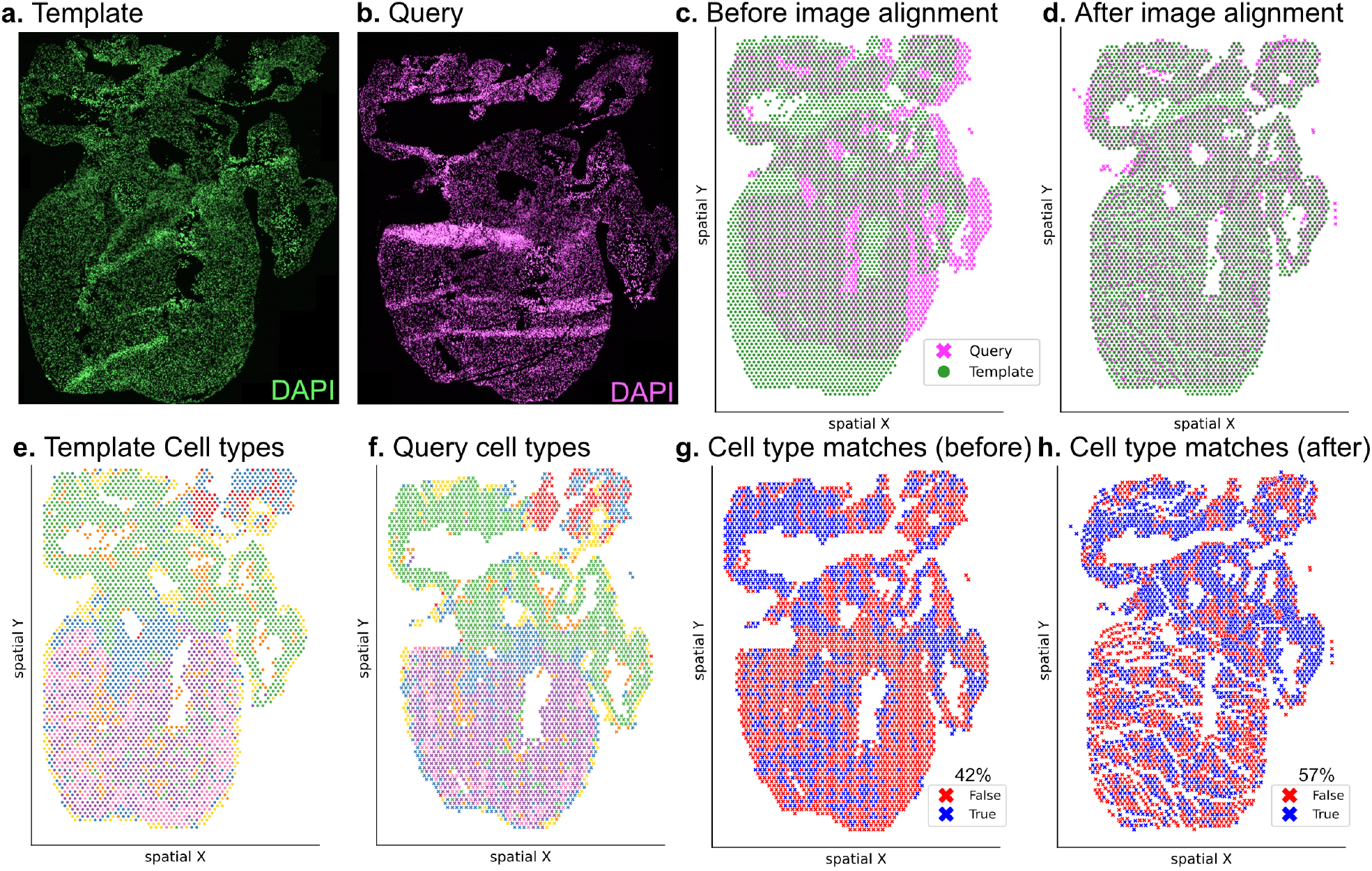
ISS heart dataset overview. a. Template slide showing pseudo-colored nuclear (DAPI, green) stain. b. Query near-serial section slide showing pseudo-colored nuclear (DAPI, pink) stain. c. Overlay of the two samples before image alignment. d. Overlay of the two slices after image-based alignment using Valis. e. Spatial scatterplot showing cell types in pseudospots generated on the ISS data on template and f. query slides. g. Cell type matches before alignment (42% match). h. Cell type matches after image alignment (57% match, ‘Initial’ in Fig. 3c-d).

The overall computational complexity of SAME is characterized by an 𝒪 (*nk*^3^) setup phase followed by an exponential worst-case solving phase. The core optimization problem is formulated as a 0-1 integer linear program (ILP), which belongs to the class of *NP*-hard problems. However, practical performance depends critically on the problem structure and the efficiency of Gurobi’s branch-and-bound implementation, which often achieves near-optimal solutions within reasonable time limits for real biological datasets. The parameter *k* serves as a crucial tuning mechanism, as reducing *k* from the theoretical maximum of |*T*| (all template cells) to practical values (typically 6-8) dramatically reduces both space and time complexity while maintaining biological relevance through local spatial matching (Fig. S2). Additionally, for a fixed *k*, the ‘penalty’ parameter that controls triangle orientation violations substantially influences solving time: higher penalty values create more tightly constrained optimization problems that require longer computation times as the solver must work harder to find feasible solutions that preserve spatial topology, while lower penalty values allow more triangle violations but enable faster convergence at the cost of reduced spatial coherence (Fig. S3).

**Figure S2.**
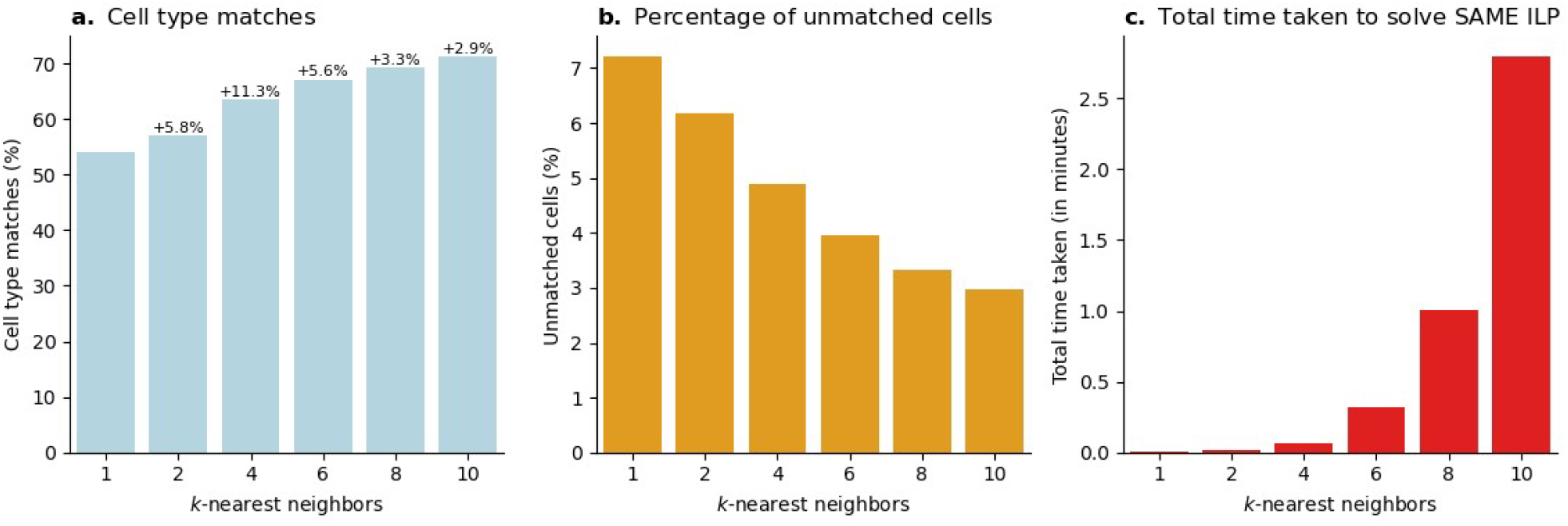
Analysis of kNN parameter on SAME for triangle penalty=5. a. Comparison of cell type matches (as a % of total query cells) with kNN parameter showing diminishing returns with increase in k above kNN value of 4. b. Change in percentage of unmatched cells with kNN parameter. c. Change in total time taken to solve SAME ILP with kNN parameter.

**Figure S3.**
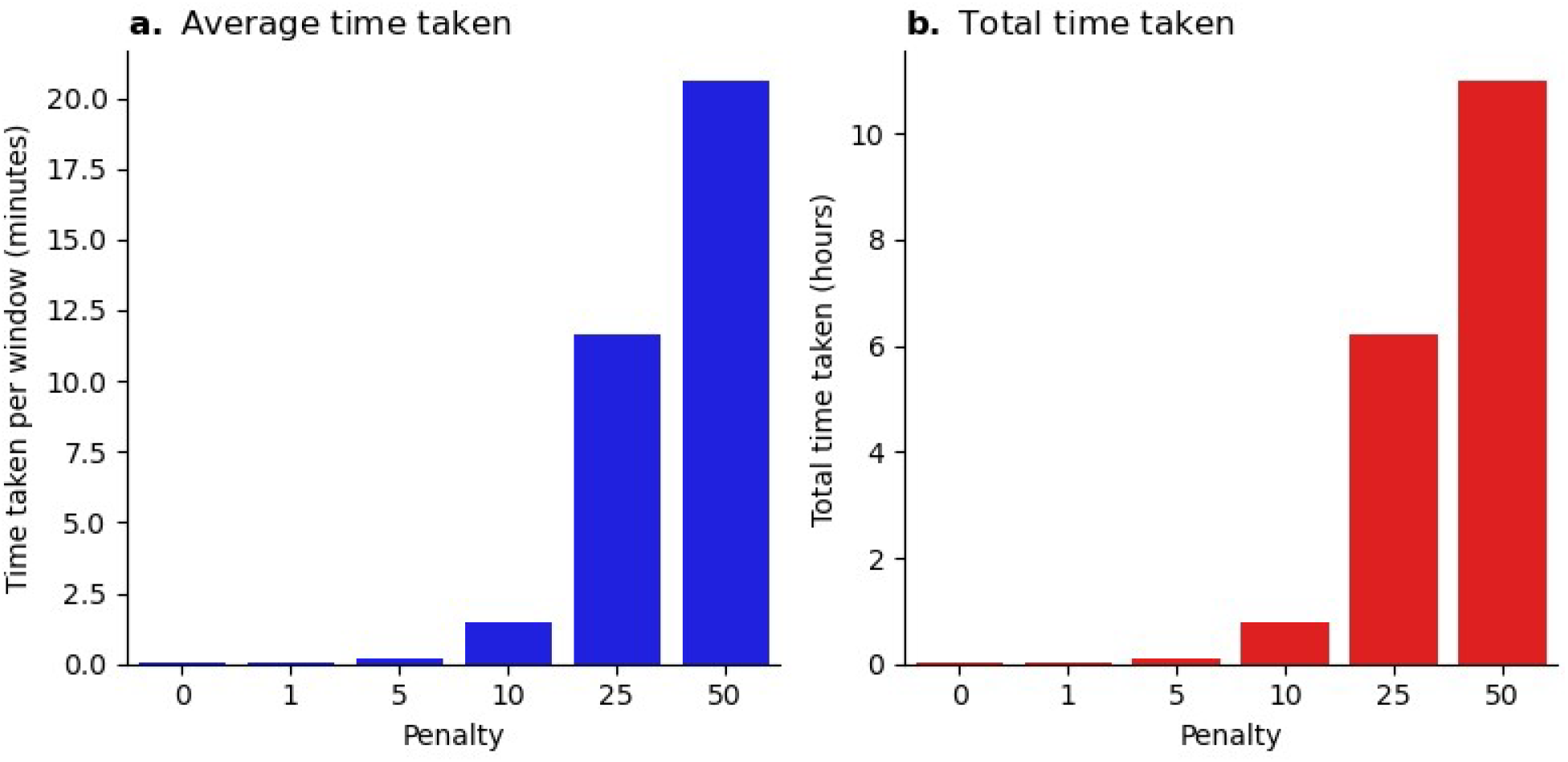
Effect of triangle penalty on SAME runtime for ISS Heart dataset (kNN=8). a. Average run time taken per sliding window. b. Total time taken to solve SAME ILP.

### Note S2: Parameter analysis for SAME

We systematically evaluated how SAME’s key parameters affect both computational performance and alignment quality for ISS heart dataset. The *k*-nearest neighbor parameter, which defines the search space for potential cell matches, showed a critical trade-off between accuracy and runtime. When we varied *k* from 2 to 10 neighbors on the ISS heart dataset for a fixed triangle penalty parameter (penalty=5), cell type matching accuracy improved from 53.9% to 71.3%, with most gains achieved around *k* = 4 (Fig. S2a). Importantly, increasing *k* dramatically reduces the fraction of unmatched cells from 7.2% at *k* = 2 to 2.9% at *k* = 10 (Fig. S2b), suggesting that larger search neighborhoods help resolve ambiguous phenotype matches in regions with mixed cell populations. However, computational cost scaled polynomially with *k*, increasing from a few seconds to 2.8 minutes for the complete ILP solution on a cloud computational infrastructure with 40 vCPUs and 64GB RAM (Fig. S2c). This scaling behavior reflects the 𝒪 (*nk*^3^) complexity of constraint generation, where each additional neighbor exponentially expands the space of possible triangle-to-triangle correspondences.

To understand how spatial coherence constraints affect SAME’s behavior, we analyzed the triangle penalty parameter across values from 0 to 50. This parameter weights the cost of triangle orientation inversions in the ILP objective function. With penalty=0, SAME solved rapidly (average 0.04 minutes per sliding window) but permitted triangle violations throughout the tissue (Fig. S3a, S4a). As we increased the penalty, runtime grew exponentially, reaching 20.4 minutes per window at penalty=50 (Fig. S3a-b). This computational burden arises because the ILP solver must explore increasingly constrained solution spaces to find feasible alignments that preserve spatial topology.

Spatial analysis of triangle violations revealed how the penalty parameter controls the location and extent of space-tearing transforms. At penalty=0, 18.3% of triangles underwent orientation inversion, distributed across the entire tissue (Fig. S4a). Increasing penalty to 5 reduced violations to 35%, primarily concentrated at tissue boundaries and regions of high cellular density (Fig. S4c). At penalty=50, only 4.4% of triangles violated orientation constraints, restricted to unavoidable topological mismatches between serial sections (Fig. S4l). Remarkably, cell type matching accuracy remained robust across penalty values, decreasing only from 73.1% to 71.2% as penalty increased from 0 to 50 (Fig 3c, Fig. S4d-f, j-l). These results establish that SAME can maintain high biological correspondence while providing user-controlled flexibility over spatial distortions.

**Figure S4.**
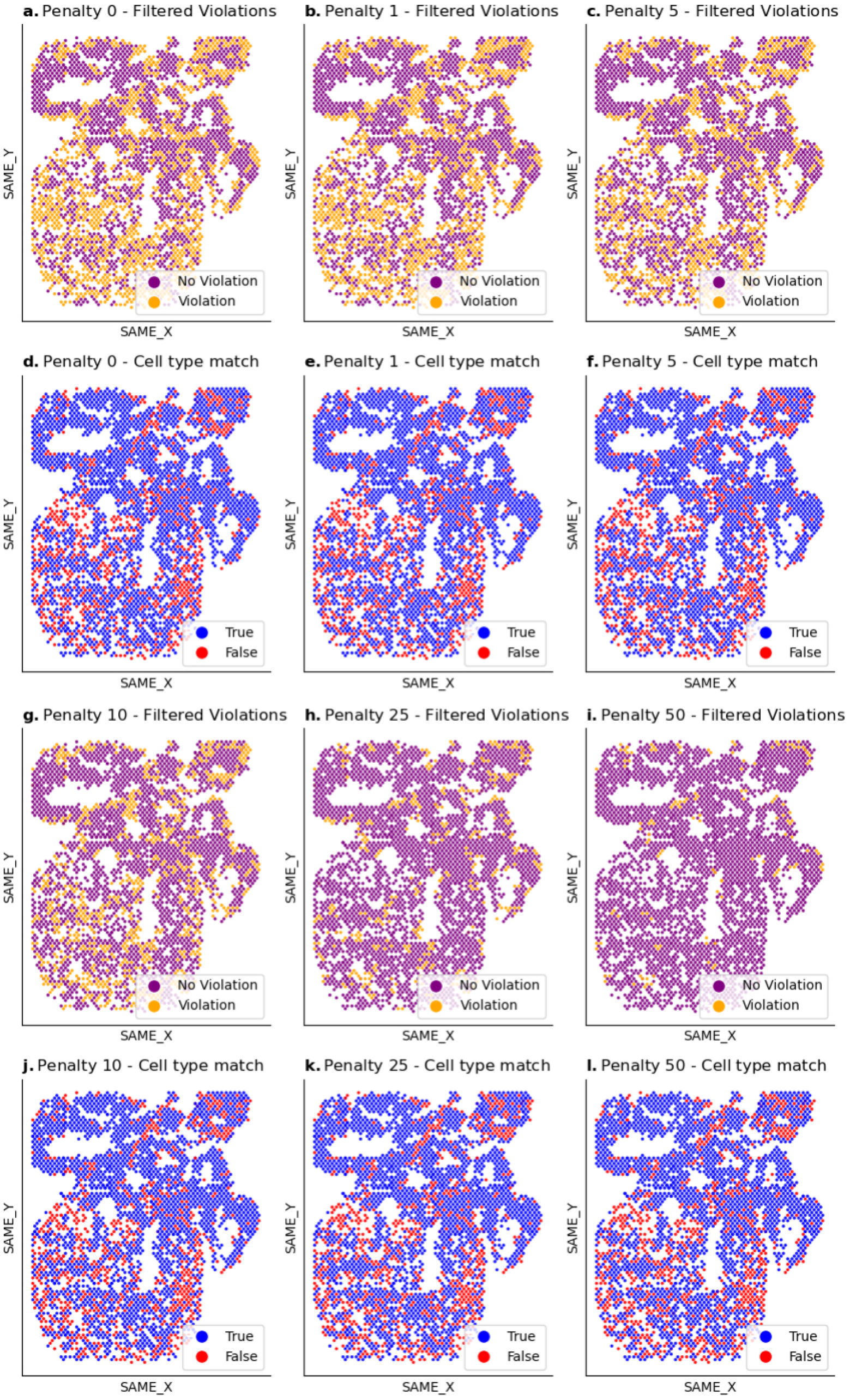
Spatial plots showing the relationship between triangle violations and cell type matches across triangle penalty parameters.

**Figure S5.**
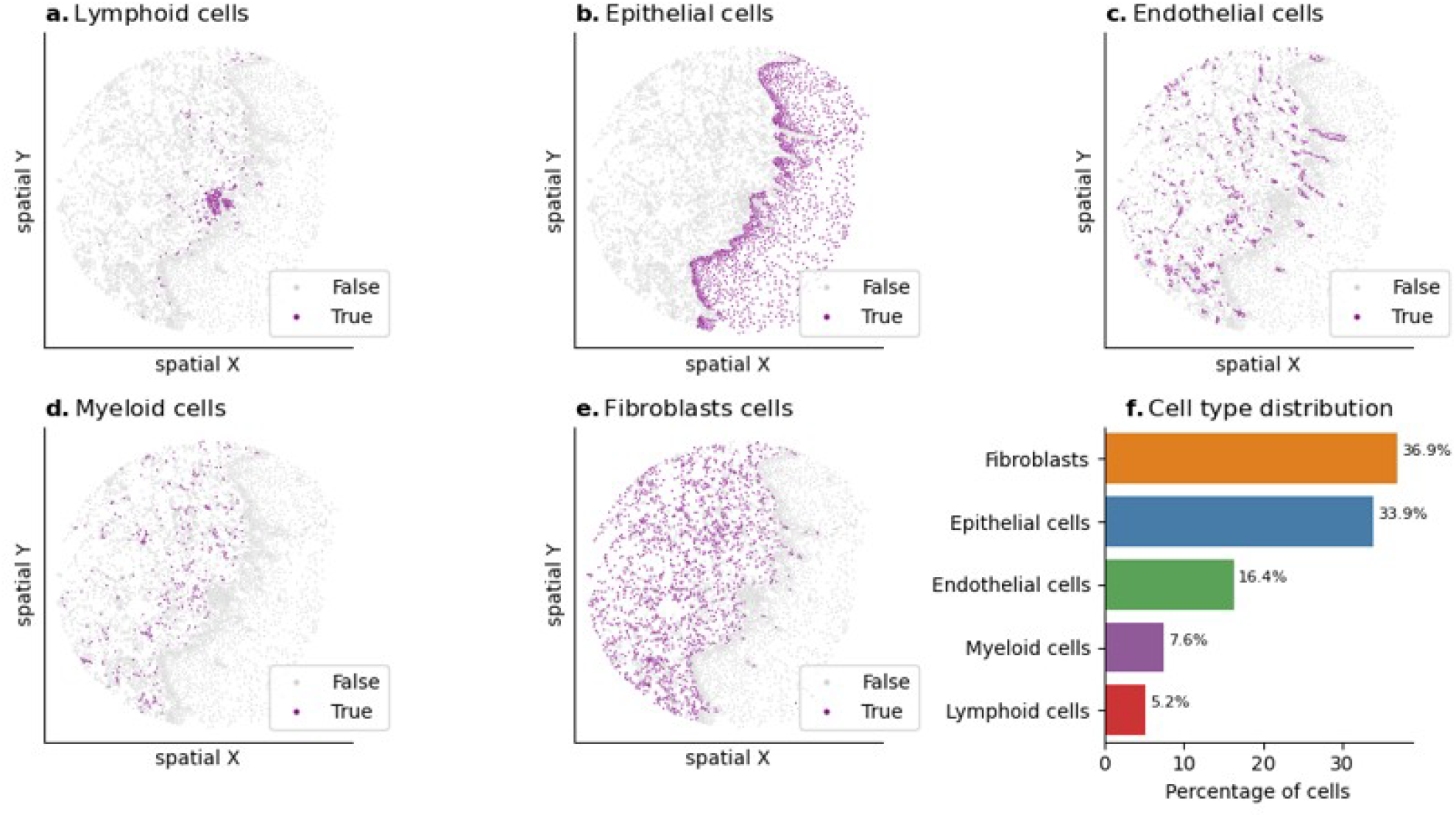
a-e. Spatial plots showing the cell type distribution from an unsupervised clustering of protein expression on the PCF query slide from dorsal tongue. f. Cell type distribution in the PCF slide.

**Figure S6.**
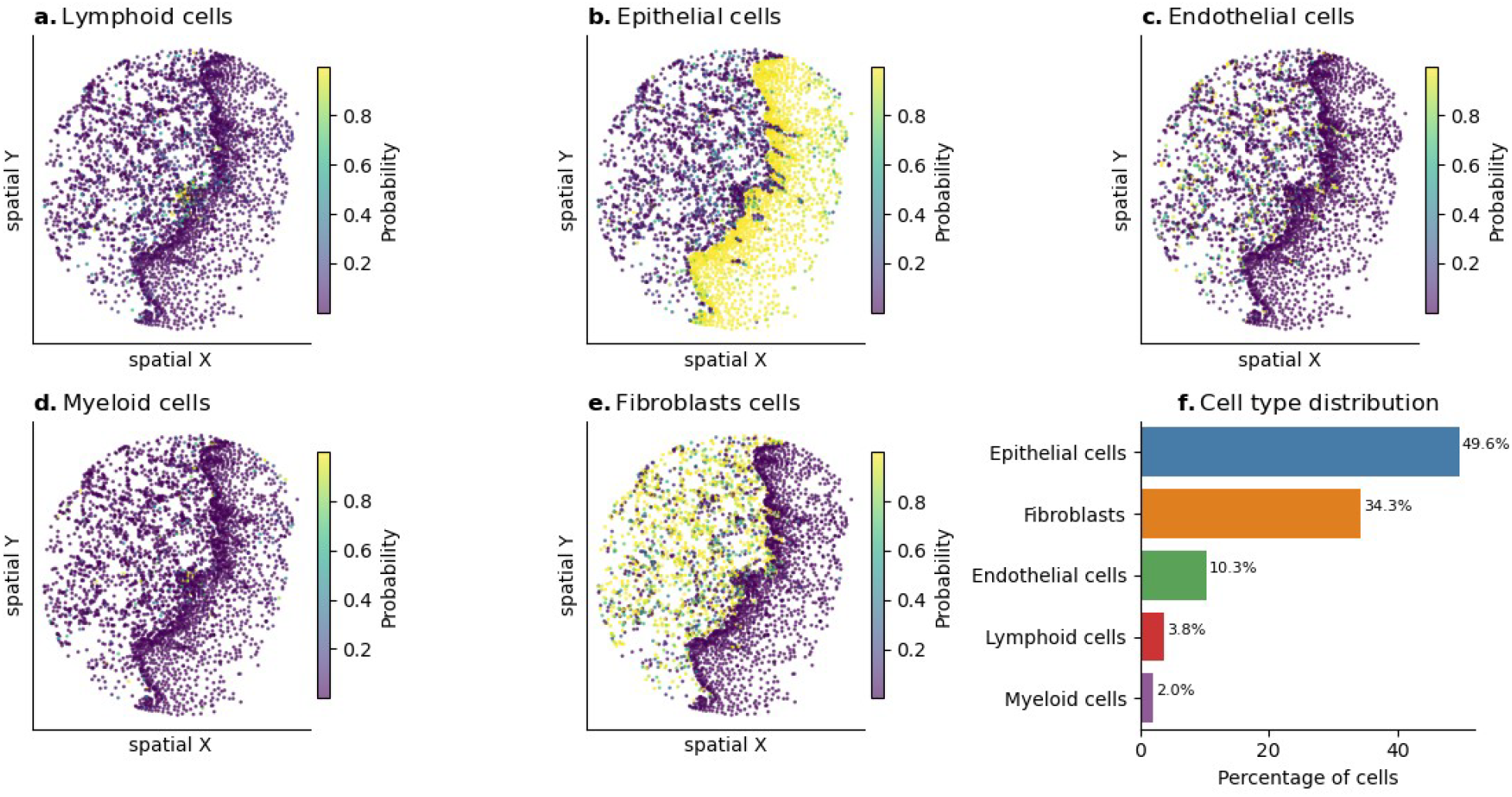
a-e. Spatial plots showing the maximum cell type probability distribution from a logistic regression classifier for mRNA counts on the MERFISH template slide from dorsal tongue. f. Cell type distribution in the MERFISH slide.

**Figure S7.**
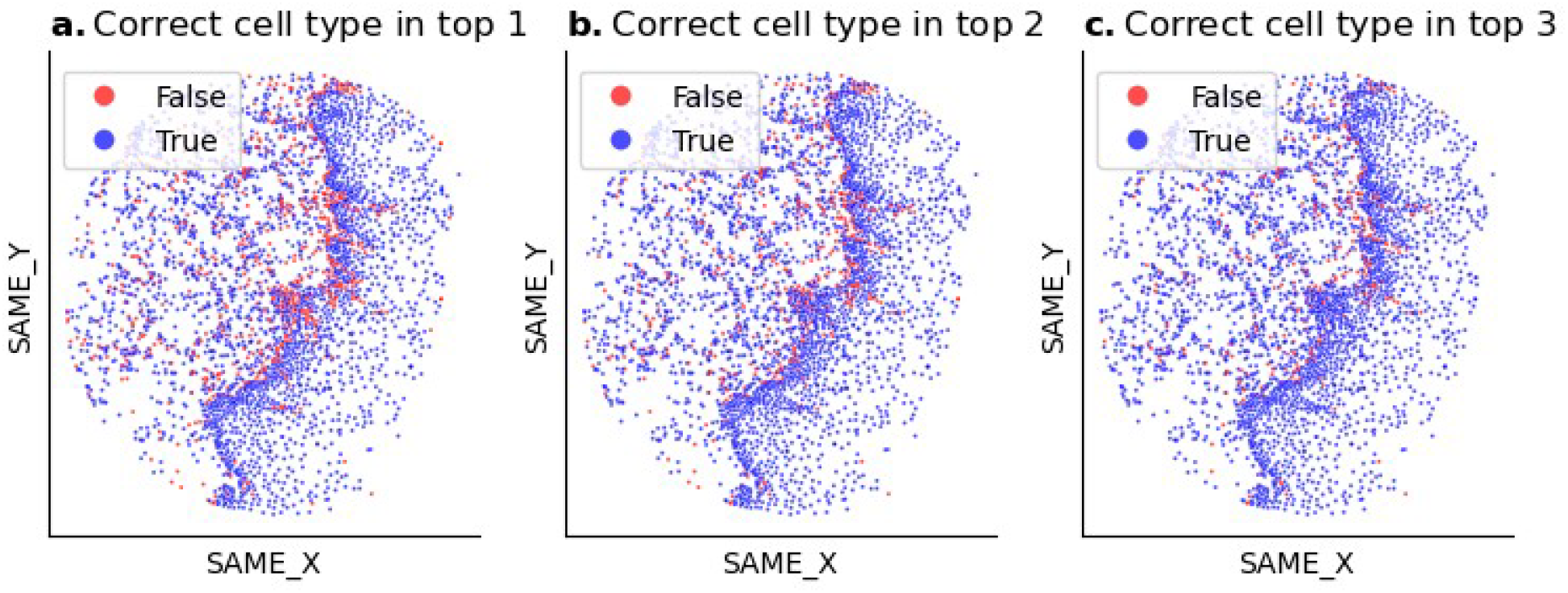
SAME cell-type matches are robust to low confidence annotation due to MERSCOPE’s noisy RNA expression levels in dorsal tongue sample. When SAME cell-type annotations differed between modalities, we assessed whether this was due to lower confidence in one modality. a. PCF cell types match the top predicted cell type in MERFISH (78.5%). b. PCF cell types appear within the top 2 predicted cell types in MERFISH (85.1%). c. PCF cell types appear within the top 3 predicted cell types in MERFISH (89.9%). Blue points indicate successful matches; red points indicate no match within the specified ranking.

**Figure S8.**
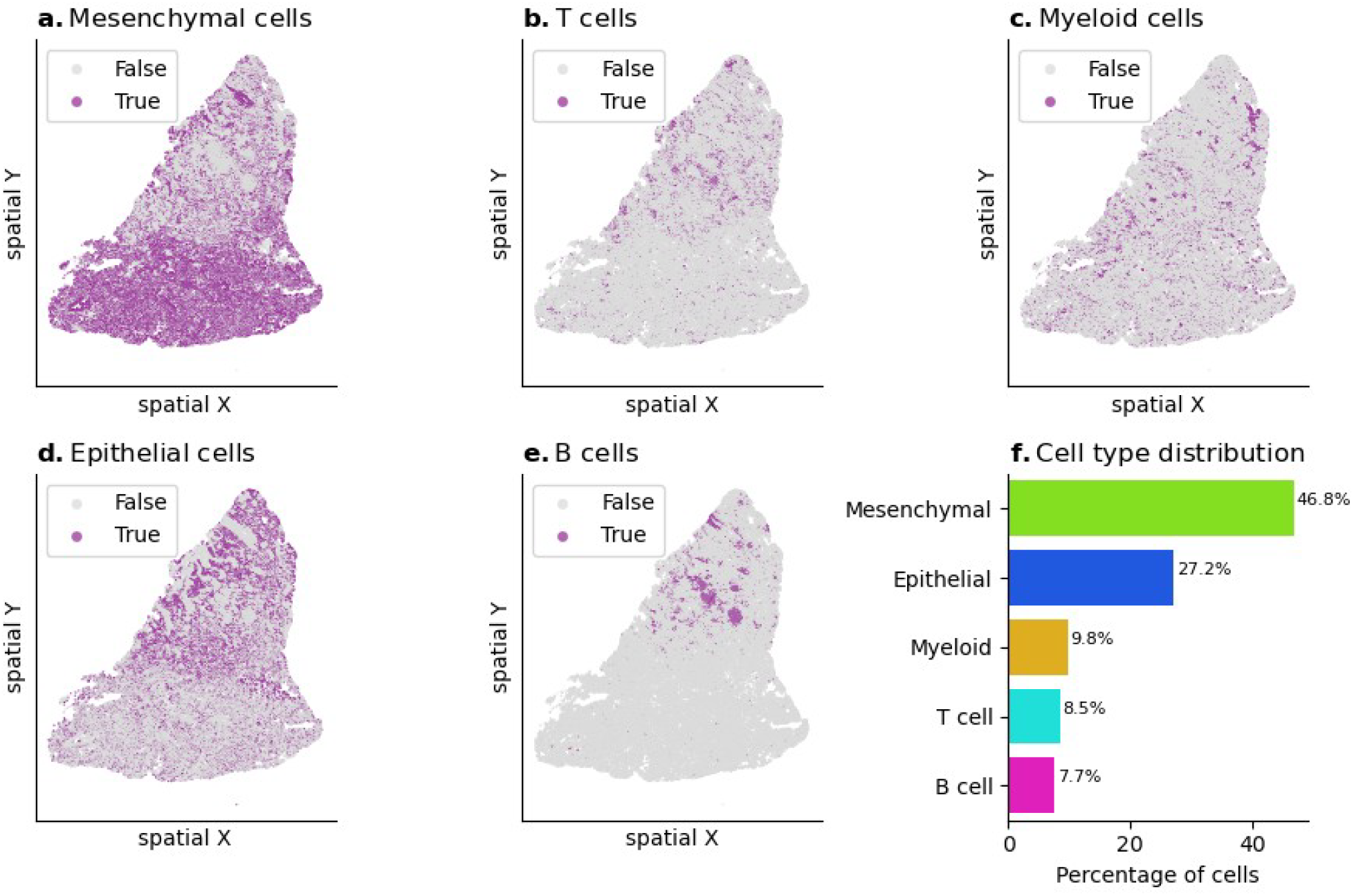
a-e. Spatial plots showing the cell type distribution from an unsupervised clustering of protein expression on the PCF query slide from lung adenocarcinoma. f. Cell type distribution in the PCF slide.

**Figure S9.**
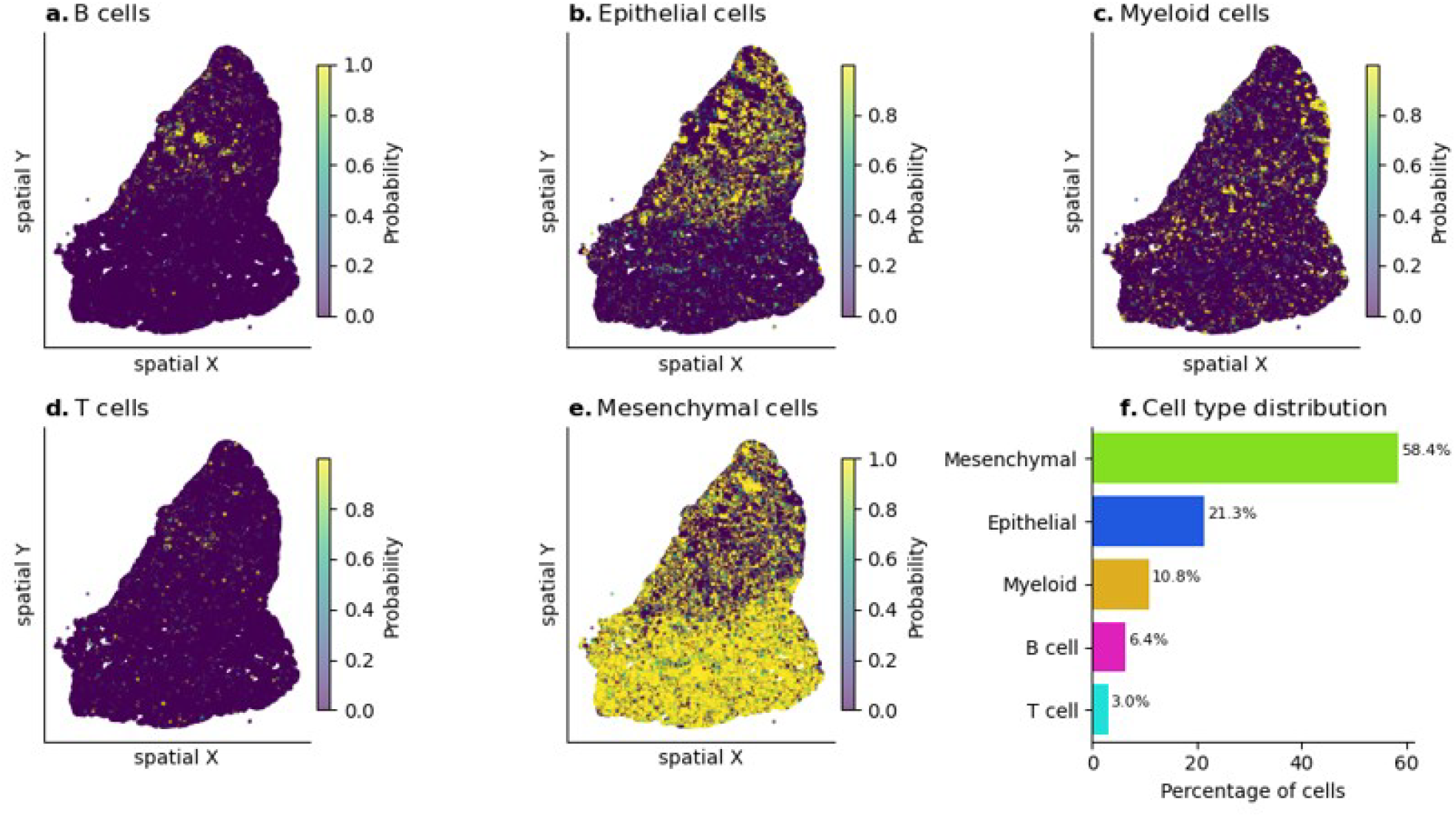
a-e. Spatial plots showing the maximum cell type probability distribution from a logistic regression classifier for mRNA counts on the Xenium template slide from lung adenocarcinoma. f. Cell type distribution in the Xenium slide.

**Figure S10.**
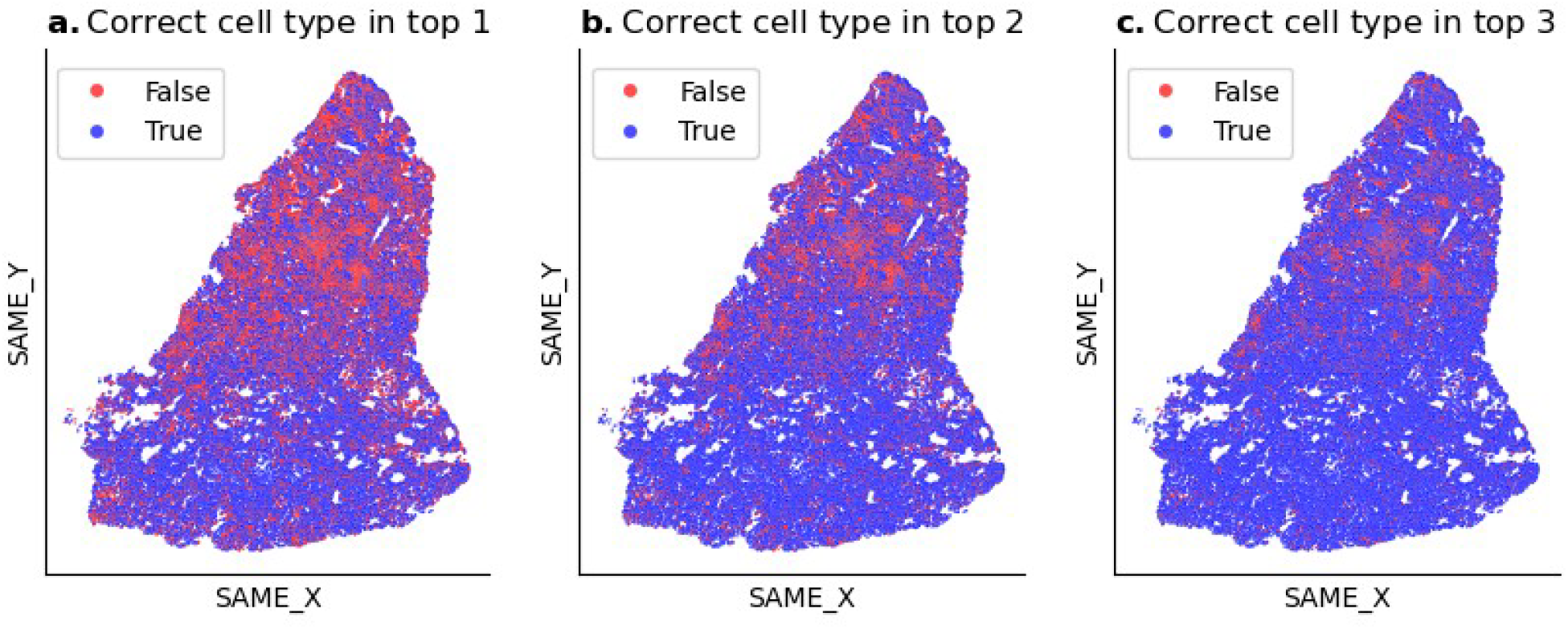
SAME cell-type matches are robust to low confidence annotations due to Xenium’s noisy RNA expression levels in lung adenocarcinoma sample. When SAME cell-type annotations differed between modalities, we assessed whether this was due to lower confidence in one modality. a. PCF cell types match the top predicted cell type in Xenium (63.4%). b. PCF cell types appear within the top 2 predicted cell types in Xenium (76.1%). c. PCF cell types appear within the top 3 predicted cell types in Xenium (85.6%). Blue points indicate successful matches; red points indicate no match within the specified ranking.

**Figure S11.**
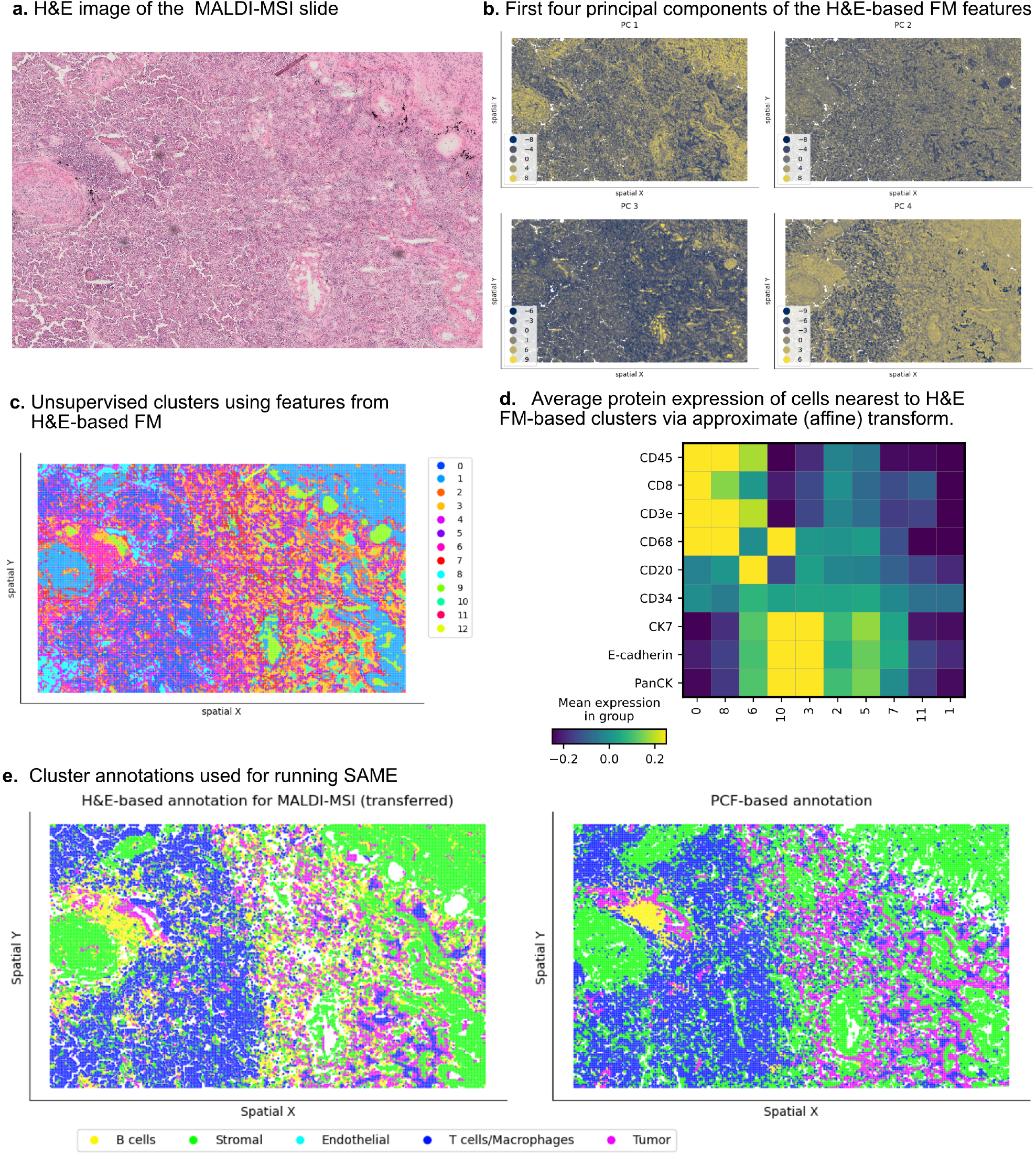
Preprocessing H&E-based staining for MALDI-MSI imaging slide. a. H&E image of the MALDI-MSI slide. b. First four principal components of the H&E-based foundation model (FM) features showing spatial variation across the tissue. c. Unsupervised clusters using features from H&E-based FM, with 13 distinct clusters identified (clusters 4,9, and 12 correspond to background; not shown). d. Average protein expression of cells spatially closest to H&E FM-based clusters via approximate (affine) transform, showing cluster-specific protein expression patterns. e. Spatial plots showing final cluster labels used for running SAME-based alignment on two serial sections

**Table S1.**
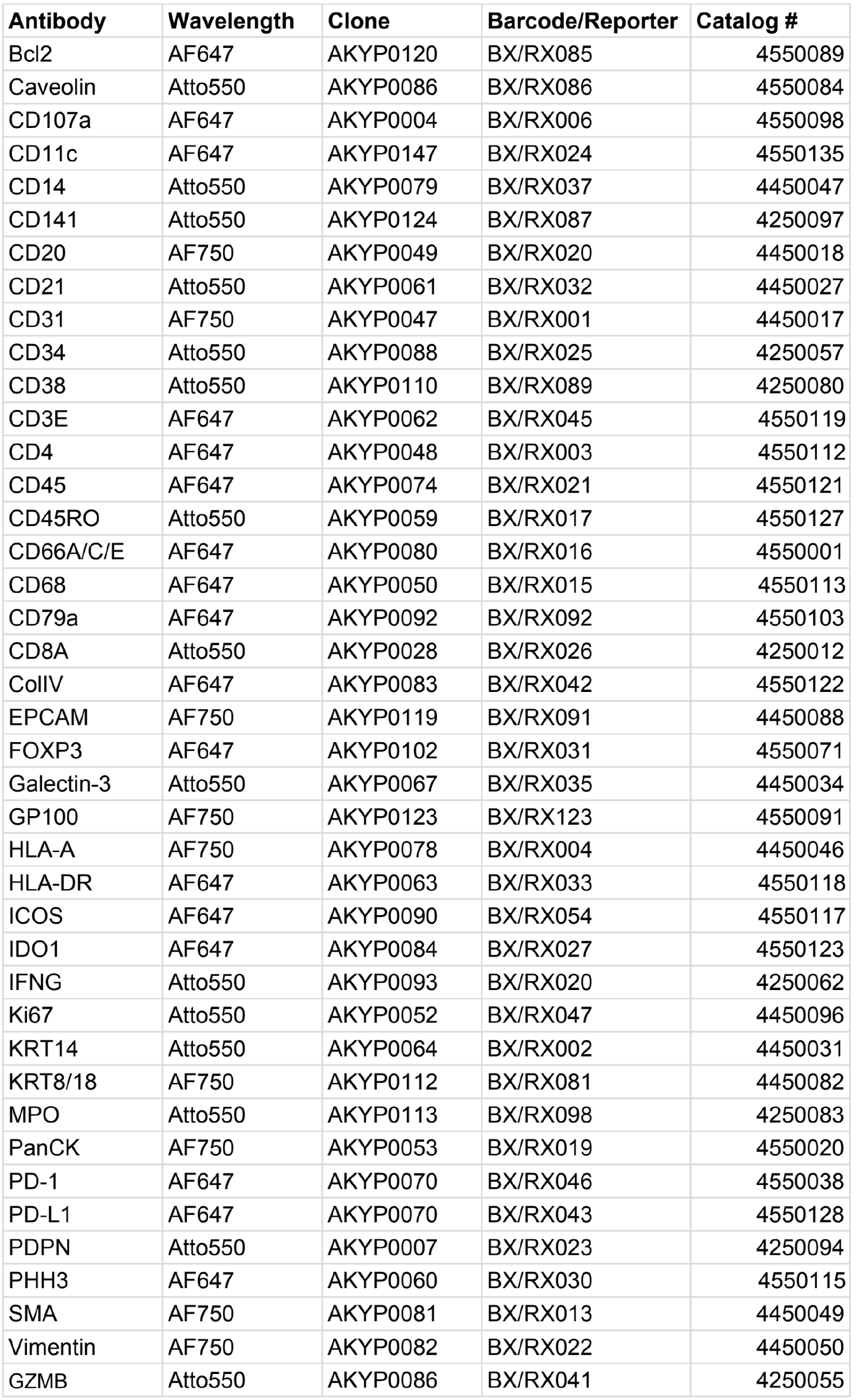
Phenocycler-Fusion (PCF) antibody panel used for dorsal tongue sample.

**Table S2.**
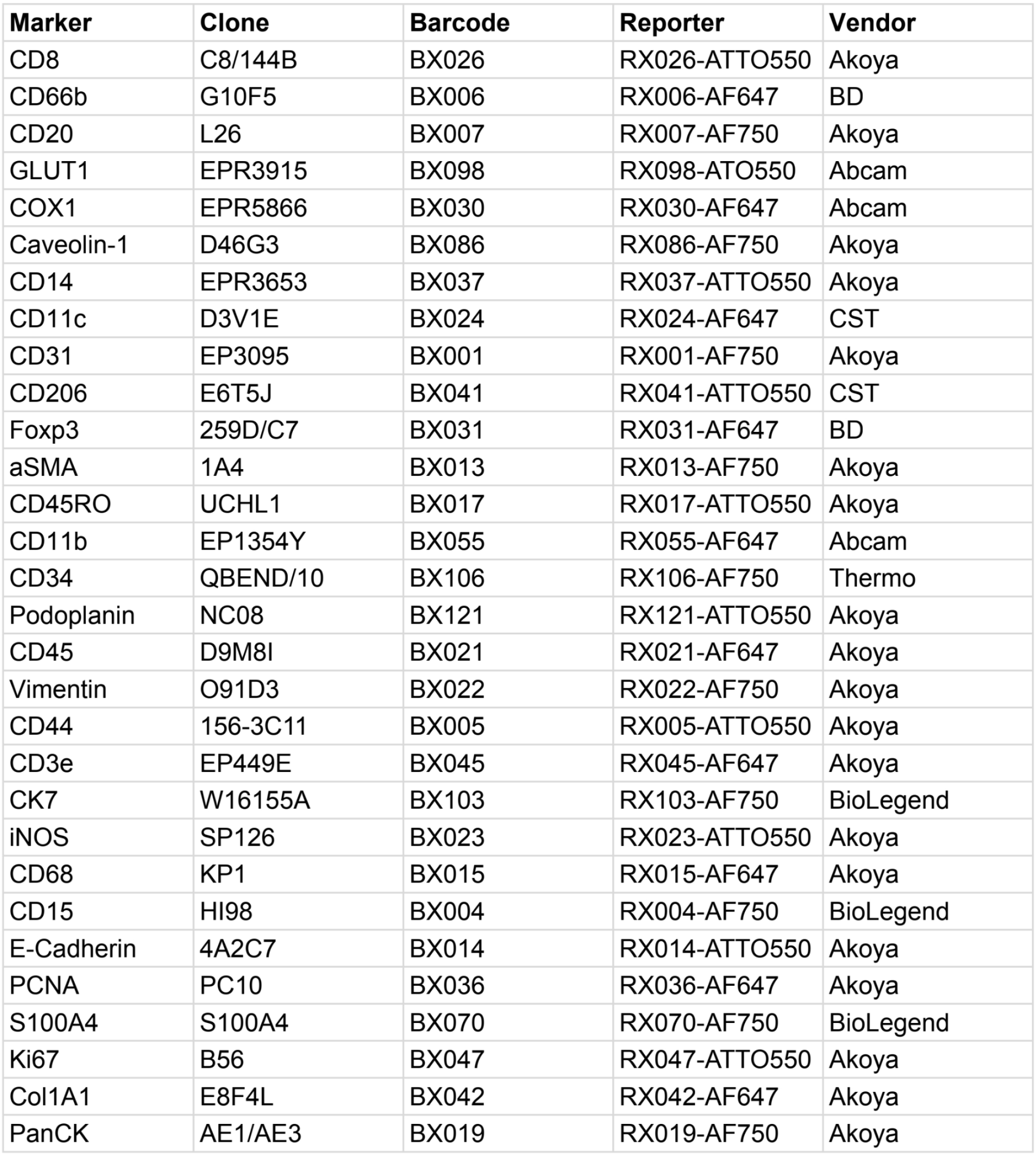
PCF antibody panel used for lung adenocarcinoma sample.

| **Marker** | **Clone** | **Barcode** | **Reporter** | **Vendor** |
| --- | --- | --- | --- | --- |
| CD8 | C8/144B | BX026 | RX026-ATTO550 | Akoya |
| CD66b | G10F5 | BX006 | RX006-AF647 | BD |
| CD20 | L26 | BX007 | RX007-AF750 | Akoya |
| GLUT1 | EPR3915 | BX098 | RX098-ATO550 | Abcam |
| COX1 | EPR5866 | BX030 | RX030-AF647 | Abcam |
| Caveolin-1 | D46G3 | BX086 | RX086-AF750 | Akoya |
| CD14 | EPR3653 | BX037 | RX037-ATTO550 | Akoya |
| CD11c | D3V1E | BX024 | RX024-AF647 | CST |
| CD31 | EP3095 | BX001 | RX001-AF750 | Akoya |
| CD206 | E6T5J | BX041 | RX041-ATTO550 | CST |
| Foxp3 | 259D/C7 | BX031 | RX031-AF647 | BD |
| aSMA | 1A4 | BX013 | RX013-AF750 | Akoya |
| CD45RO | UCHL1 | BX017 | RX017-ATTO550 | Akoya |
| CD11b | EP1354Y | BX055 | RX055-AF647 | Abcam |
| CD34 | QBEND/10 | BX106 | RX106-AF750 | Thermo |
| Podoplanin | NC08 | BX121 | RX121-ATTO550 | Akoya |
| CD45 | D9M8I | BX021 | RX021-AF647 | Akoya |
| Vimentin | O91D3 | BX022 | RX022-AF750 | Akoya |
| CD44 | 156-3C11 | BX005 | RX005-ATTO550 | Akoya |
| CD3e | EP449E | BX045 | RX045-AF647 | Akoya |
| CK7 | W16155A | BX103 | RX103-AF750 | BioLegend |
| iNOS | SP126 | BX023 | RX023-ATTO550 | Akoya |
| CD68 | KP1 | BX015 | RX015-AF647 | Akoya |
| CD15 | HI98 | BX004 | RX004-AF750 | BioLegend |
| E-Cadherin | 4A2C7 | BX014 | RX014-ATTO550 | Akoya |
| PCNA | PC10 | BX036 | RX036-AF647 | Akoya |
| S100A4 | S100A4 | BX070 | RX070-AF750 | BioLegend |
| Ki67 | B56 | BX047 | RX047-ATTO550 | Akoya |
| Col1A1 | E8F4L | BX042 | RX042-AF647 | Akoya |
| PanCK | AE1/AE3 | BX019 | RX019-AF750 | Akoya |

## Footnotes

† We avoid the term ‘non-diffeomorphic’ as its mathematical connotation also covers topology-preserving transforms that are simply non-bijective or non-smooth

